# *Neuroplastin* genetically interacts with *Cadherin 23* and the encoded isoform Np55 is sufficient for cochlear hair cell function and hearing

**DOI:** 10.1101/2021.11.10.468016

**Authors:** Sherylanne Newton, Fanbo Kong, Adam J Carlton, Carlos Aguilar, Andrew Parker, Gemma F Codner, Lydia Teboul, Sara Wells, Steve DM Brown, Walter Marcotti, Michael R Bowl

## Abstract

Mammalian hearing involves the mechanoelectrical transduction (MET) of sound-induced fluid waves in the cochlea. Essential to this process are the specialised sensory cochlear cells, the inner (IHCs) and outer hair cells (OHCs). While genetic hearing loss is highly heterogeneous, understanding the requirement of each gene will lead to a better understanding of the molecular basis of hearing and also to therapeutic opportunities for deafness. The *Neuroplastin* (*Nptn*) gene, which encodes two protein isoforms Np55 and Np65, is required for hearing, and homozygous loss-of-function mutations that affect both isoforms lead to profound deafness in mice. Here we have utilised several distinct mouse models to elaborate upon the spatial, temporal, and functional requirement of *Nptn* for hearing. While we demonstrate that both Np55 and Np65 are present in cochlear cells, characterisation of a Np65-specific mouse knockout shows normal hearing thresholds indicating that Np65 is functionally redundant for hearing. In contrast, we find that *Nptn*-knockout mice have significantly reduced maximal MET currents and MET channel open probabilities in mature OHCs, with both OHCs and IHCs also failing to develop fully mature basolateral currents. Furthermore, comparing the hearing thresholds and IHC synapse structure of *Nptn*-knockout mice with those of mice that lack *Nptn* only in IHCs and OHCs shows that the majority of the auditory deficit is explained by hair cell dysfunction, with abnormal afferent synapses contributing only a small proportion of the hearing loss. Finally, we show that continued expression of NEUROPLASTIN in OHCs of adult mice is required for membrane localisation of Plasma Membrane Ca^2+^ ATPase 2 (PMCA2), which is essential for hearing function. Moreover, *Nptn* haploinsufficiency phenocopies *Atp2b2* (encodes PMCA2) mutations, with heterozygous *Nptn*-knockout mice exhibiting hearing loss through genetic interaction with the *Cdh23^ahl^* allele. Together, our findings provide further insight to the functional requirement of *Neuroplastin* for mammalian hearing.

**Author Summary:** Sensorineural hearing loss, caused by problems with sensory cells in the cochlea or the auditory nerve, is the most common type of hearing loss. Mutations in *Neuroplastin* have already been implicated in deafness in mice. We have used mutant mouse models to investigate where *Neuroplastin* is expressed in the cochlea and its function. When mice do not express a functioning copy of *Neuroplastin* they have disruptions to the primary sensory synapse. We show that although synaptic disruption contributes to the loss of hearing function it is not the primary cause. Instead, continued expression of *Neuroplastin* is needed to maintain the localisation of Plasma Membrane Ca^2+^ ATPase 2 channels which help regulate calcium flow. We have also shown that two types of NEUROPLASTIN protein (isoforms) are both expressed within the cochlea, although only one of these isoforms needs to be expressed for normal hearing. Finally, we also demonstrate that the hearing loss caused by the absence of *Neuroplastin* is made worse when combined with a common mutation within a gene called *Cadherin 23* (*Cdh23^ahl^*). This is an important finding as although there are currently no human patients with an identified *NEUROPLASTIN* mutation, it may be involved in human deafness in combination with other mutations.

## Introduction

The mammalian cochlea is an extremely complex and organised structure consisting of multiple cell types that act in concert to convert sound into neuronal signals. In particular, within the organ of Corti are two functionally distinct sensory cells: the inner hair cells (IHCs), which relay sound stimuli to the brain via the release of glutamate from ribbon synapses onto type I spiral ganglion neurons; and the outer hair cells (OHCs) which mechanically amplify sound stimuli through the generation of voltage-dependent axial forces on the organ of Corti. The function of both IHCs and OHCs is driven by mechanoelectrical transduction (MET) channels located at the tips of modified microvilli called stereocilia at the apex of each cell, activated following deflection of the stereocilia bundle. Though we now have a good understanding of the specific function of auditory hair cells, it is still largely unknown how the functional development of these complex sensory cells is orchestrated at a molecular level. Calcium ions (Ca^2+^) have several essential roles in the cochlea, including contribution to the total MET current [1], and driving adaptation of the MET channels, reducing their open probability [2]. Highly coordinated Ca^2+^-signaling is also required for the maturation of afferent synapses on both IHCs [3] and OHCs [4], and in mature IHCs, the influx of Ca^2+^, primarily through Ca_V_1.3 channels located at each active zone, facilitates the release of vesicles onto the afferent terminals.

NEUROPLASTIN (ENSMUSG00000032336), together with BASIGIN (ENSMUSG00000023175) and EMBIGIN (ENSMUSG00000021728), comprise a small family of neural cell adhesion molecules (NCAM), which are an integral component of the synaptic membrane and are proposed to mediate cellular processes such as synaptic plasticity and neuronal differentiation [5–9]. Furthermore, the study of mouse mutants has shown a role for this family in sensory function, including vision and hearing [10, 11]. Of these, the role of NEUROPLASTIN in hearing is the most studied, with loss-of-function mutations causing profound early-onset hearing loss [12, 13]. However, there are many outstanding questions regarding the role and requirement of NEUROPLASTIN for mammalian hearing. The *Neuroplastin* gene (*Nptn*) encodes two protein isoforms: the larger Np65 that consists of three extracellular Ig domains (Ig1-3), a transmembrane domain and a short intracellular C-terminal; and, Np55 in which exon 2 is skipped to produce a shorter NEUROPLASTIN without Ig1 [14]. Np55 is reported to be localised to outer hair cell (OHC), but not inner hair cell (IHC) stereocilia, where it is required for their correct coupling to the tectorial membrane [13]. Localisation of NEUROPLASTIN has also been reported at the basolateral membrane of IHCs, where it plays a critical role in the formation of mature ribbon synapses, which was hypothesised to involve Np65 [12]. To date, all of the *Nptn* alleles studied in relation to hearing involve loss of both Np55 and Np65 [12, 13, 15].

Here we have studied several *Neuroplastin* knockout mouse models to further elaborate upon the role of this NCAM in mammalian hearing. We find that deletion of Np65 alone does not cause deafness, thereby demonstrating that Np55 is sufficient to support auditory development and function. Moreover, we demonstrate that while IHC synaptopathy is a significant pathological feature, synapse formation does not require Np65, nor is it the main driver of hearing loss in *Nptn*-null mice. Furthermore, continued expression of *Neuroplastin* in adulthood is required to maintain hearing function. Finally, we demonstrate that the expressivity of the *Nptn* loss-of-function auditory phenotype is potentiated by strain background, showing a genetic interaction between *Nptn* and the strain-specific *Cadherin 23* (Otocadherin, ENSMUSG00000012819) mutant allele, *Cdh23^ahl^,* that is present in C57BL/6 mice. This effect is likely mediated through the NEUROPLASTIN-dependent localisation of PMCA2 channels in stereocilia.

## Results

### Both Np65 and Np55 are cochlear expressed and absent in the *Nptn^tm1b^* deafness mutant

To gain insight into which NEUROPLASTIN protein isoforms are required for hearing, an RT-PCR study was undertaken to determine the *Nptn* transcripts present in wild type whole cochleae. Transcripts relating to the two main protein isoforms, Np65 (*Nptn-212,* ENSMUST00000177292, exons 1 - 8, 397aa) and Np55 (*Nptn-201,* ENSMUST00000085651, exon 2 skipped, 281aa), were amplified. These differ by the presence or absence of exon 2 that encodes Ig1, respectively (Fig 1A). To assess the presence of both Np65 and Np55 isoforms in mouse cochlear tissue, Western blotting was undertaken utilising two commercially-available anti-NEUROPLASTIN antibodies: an anti-pan-Np antibody that detects both Np65 and Np55, and an anti-Np65-specific antibody (Fig 2A). For negative control lysates, we utilised the *Nptn^tm1b^* mouse mutant generated as part of the International Mouse Phenotyping Consortium (IMPC) programme, in which both isoforms are disrupted (Fig 1A, B). As NEUROPLASTIN is a highly glycosylated membrane protein, total protein lysates were first enriched for the membrane fraction before treatment with PNGase F to remove N-linked glycans [8, 12]. In the whole cochlear membrane-enriched fraction, the anti-pan-Np antibody detects bands for both Np65 and Np55 (Fig 2A), whereas the anti-Np65-specific antibody produced only a single band corresponding to Np65 (Fig 2A), confirming the presence of both Np55 and Np65 protein isoforms in wild type cochleae. However, for mutant *Nptn^tm1b/tm1b^* lysates, no bands were present when probing the membrane-enriched fractions using either antibody (Fig 2A). These results validate the specificity of the antibodies for immunoblotting and also demonstrate the absence of both NEUROPLASTIN isoforms in *Nptn^tm1b/tm1b^* cochleae. Similar results were achieved when probing brain lysates prepared from *Nptn^+/+^* and *Nptn^tm1b/tm1b^* mice (Fig 2B). Localisation of NEUROPLASTIN protein in cochleae was also assessed employing immunolabeling of cochlear cryosections. In wild type cochlear tissue, the anti-pan-Np antibody produced strong labelling of outer hair cell (OHC) stereocilia, the body of inner hair cells (IHCs) and spiral ganglion neurons (SGNs) (Fig 2C). The anti-Np65-specific antibody produced a signal in the cell body of IHCs only (Fig 2D). In addition, both antibodies labelled the non-sensory cells lateral to the OHCs, with the anti-pan-Np antibody showing diffuse membrane labelling, and the anti-Np65 antibody labelling the basal region of these cells (Fig 2C, D), which are presumed to be Hensen and Boettcher cells. Neither antibody labelled cochlear tissue derived from *Nptn^tm1b/tm1b^* mutants (Fig 2C, D). Although Np55 is preferentially localised in the stereocilia of OHCs and in SGNs, and Np65 in IHCs, our data cannot discern whether Np55 is also expressed in IHCs.

**Fig 1.**
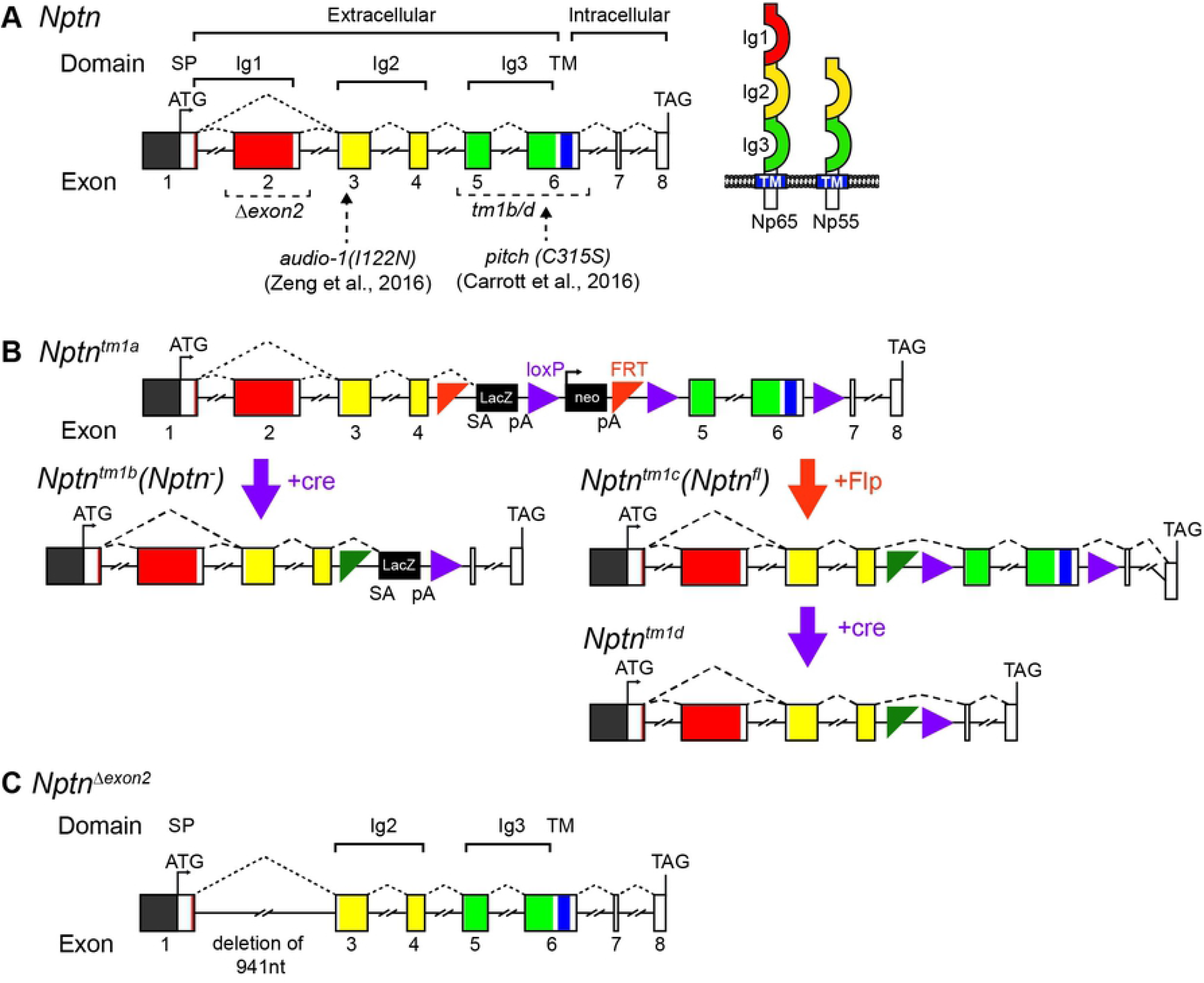
Neuroplastin gene schematic and alleles. (A) Schematic of the wild type Neuroplastin gene locus consisting of eight exons that encode: three extracellular Ig domains; a transmembrane region (TM); and, a short intracellular region. Alternative splicing involving the inclusion, or skipping, of exon 2 leads to the production of transcripts encoding the larger Np65, or smaller Np55, protein isoform, respectively. Previously reported mutant alleles *audio-1* [13] and *pitch* [12] are point mutations present in exons 3 and 6, respectively, affecting both isoforms. (B) the *Nptn^tm1a(EUCOMM)Hmgu^* (*Nptn^tm1a^*) is a knockout first, conditional-ready allele, that was used to generate both the *Nptn^tm1b(EUCOMM)Hmgu^ (Nptn^tm1b^*) knockout allele lacking exons 5 and 6, and the *Nptn^tm1c(EUCOMM)Hmgu^ (Nptn^fl^*) floxed allele. Subsequent exposure of the *Nptn^fl^* allele to cre recombinase excises exons 5 and 6, resulting in a null allele (*Nptn^tm1d^*) in cells expressing *cre.* (C) An isoform-specific Np65 knockout mutant (*Nptn^Δexon2^*) was generated using CRISPR/Cas9-mediated deletion of 941 nucleotides encompassing exon 2. Mice homozygous for the *Nptn^Δexon2^* are unable to express the larger Np65 isoform, while expression of Np55 remains.

**Fig 2.**
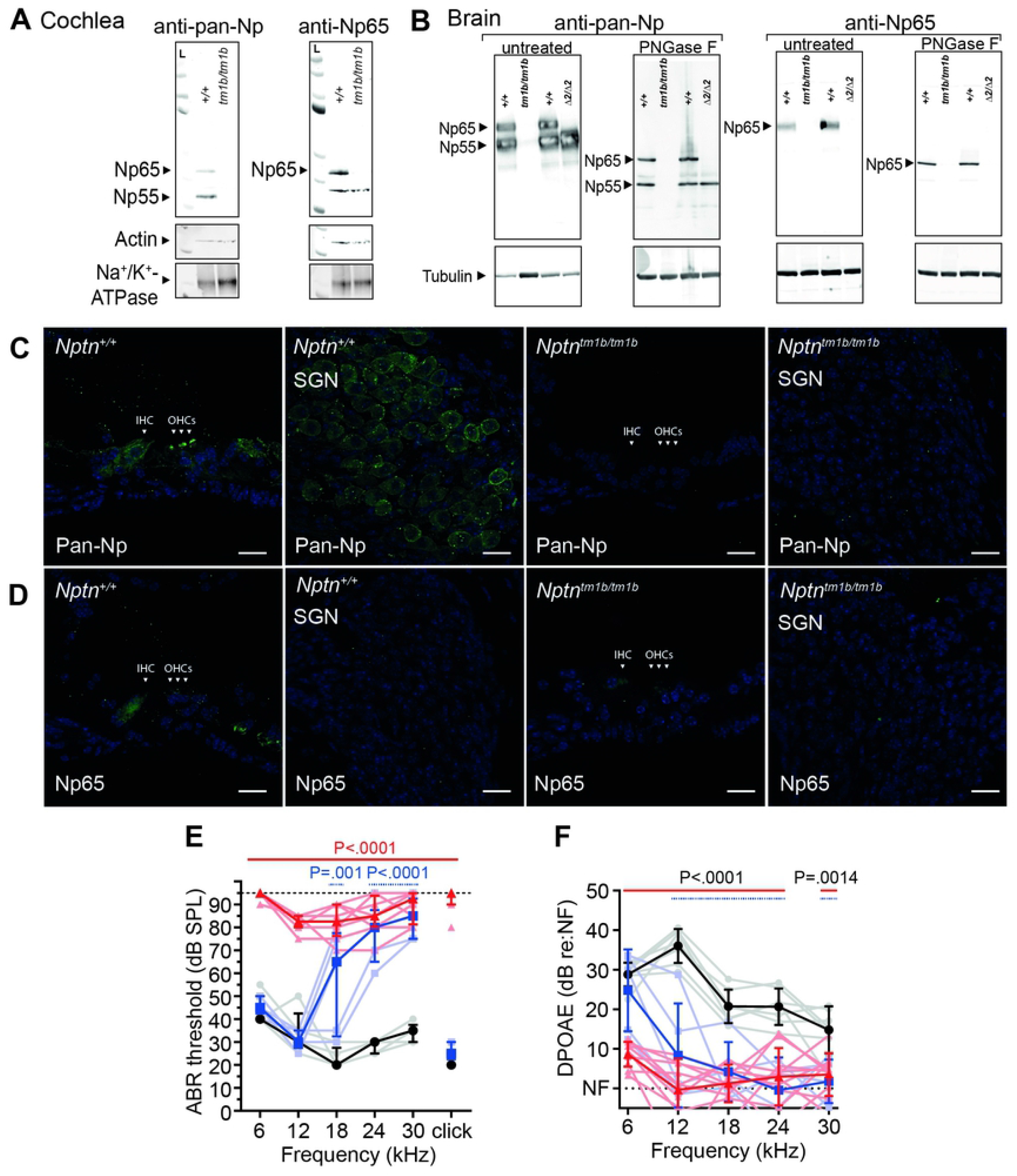
Np65 and Np55 are cochlear expressed. (A) Western blot of membrane-enriched fractions prepared from *Nptn^+/+^* and *Nptn^tm1b/tm1b^* cochleae. Lysates were treated with PNGase F to remove N-linked glycans before separation. In the wild type membrane fraction, the anti-pan-Np antibody detects bands corresponding to native Np65 (44.4 kDa) and Np55 (31.3 kDa), whereas the anti-Np65-specific antibody detects only a single band corresponding to Np65. No bands were detected in the membrane fraction prepared from *Nptn^tm1b/tm1b^* cochleae using either antibody. Na^+^/K^+^-ATPase was used as a marker protein for the plasma membrane. (B) Whole brain protein lysates prepared from *Nptn^tm1b/tm1b^ (tm1b/tm1b), Nptn^Δexon2/Δexon2^* (*Δ2/Δ2*) and their respective wild type controls, were separated by PAGE and Western blotted utilizing an anti-pan-Np antibody and an anti-Np65 antibody. For wild type lysates, both Np55 and Np65 are detected. However, neither protein isoform was detected in *Nptn^tm1b/tm1b^* lysates. Importantly, only Np55 was detected in *Nptn^Δexon2/Δexon2^* lysates, confirming the absence of Np65. Western blots were run using native (untreated) and deglycosylated (PNGase F) lysates. Tubulin is used as a loading control. (C) Cochlear immunohistochemistry of cryosections using the anti-pan-Np antibody showed labelling of outer hair cell (OHC) stereocilia, inner hair cells (IHC), support cells lateral to the hair cells, and the spiral ganglion neurons (SGN). No signal was observed in *Nptn^tm1b/tm1b^* cochlea. (D) The anti-Np65 specific antibody showed labelling of IHCs and support cells lateral to the hair cells. No signal was observed in *Nptn^tm1b/tm1b^* cochleae. Scale = 20 μm. (E) ABR threshold measures recorded from 1-month old mice showing profoundly elevated thresholds for all stimuli tested in *Nptn^tm1b/tm1b^* mice (red triangles, n = 8) compared to wild type littermates (black circle, n = 5). Heterozygous *Nptn^+/tm1b^* mice (blue squares, n = 5) exhibit a significant high frequency (≥12 kHz) hearing loss compared to control littermates. Data shown as median ± I.Q.R with individual data points. One-way ANOVA comparing to wild type, employing Dunnett’s test for multiple comparisons. (F) Average DPOAE responses recorded from 1-month old mice. Both *Nptn^tm1b/tm1b^* (n = 10) and *Nptn^+/tm1b^* (n = 5) mice have suppressed response amplitudes compared to littermate controls (n = 7). Data are mean ± SD with individual data points shown. One-way ANOVA comparing to wild type controls, employing Dunnett’s test for multiple comparisons.

IMPC-based auditory phenotyping of four *Nptn^tm1b/tm1b^* mice identified this mutation as a candidate hearing loss allele [11]. However, due to the high-throughput nature of the IMPC programme, no further study of their auditory function was undertaken. To test the consequence of the *Nptn^tm1b^* allele upon hearing function, heterozygous intercross matings were set up and the resultant offspring subject to auditory phenotyping. At four weeks of age, homozygous *Nptn^tm1b/tm1b^* mutants are profoundly deaf with ABR-threshold values ≥ 70 dB SPL across all frequencies tested, with many of these mice having no evoked ABR response up to 95 dB SPL at one or more frequencies (Fig 2E). In contrast, age-matched wild-type *Nptn^+/+^* littermate mice exhibit ABR-threshold values within the expected normal range for each of the frequencies tested (between 20 - 55 dB SPL). In contrast to previously reported *Nptn* models [12, 13], heterozygous *Nptn^+/tm1b^* littermate mice exhibit significantly elevated mean ABR-thresholds for higher frequency stimuli (≥ 18 kHz, P<0.001, one-way ANOVA) (Fig 2E).

Together, these data show that Np65 and Np55 are both present in wild type cochlear tissue, although they exhibit differential patterns of cellular expression. Furthermore, we confirm the *Nptn^tm1b^* allele is a null that causes hearing loss, which is consistent with previous studies showing that *Nptn* loss-of-function mutations cause recessive deafness [12, 13]. However, our finding that heterozygous *Nptn^+/tm1b^* mutant mice exhibit high-frequency hearing impairment is an important new finding, which impacts upon the interpretation of *NPTN* variants identified in humans with hearing loss.

### Loss of NEUROPLASTIN affects the functional maturation of OHCs

Our immunolabeling data show the localisation of Np55 in OHC stereocilia bundles (Fig. 1D). To determine if OHC activity is affected in the absence of NEUROPLASTIN we recorded distortion-product otoacoustic emissions (DPOAEs), which are an *in vivo* measure of OHC function. At 5-weeks of age, DPOAEs are significantly reduced in *Nptn^tm1b/tm1b^* mice at all frequencies tested compared to *Nptn^+/+^* littermates (Fig 2F), indicating that OHC function is likely impaired in *Nptn^tm1b/tm1b^* mutant mice. In correlation to the ABR data, *Nptn^+/tm1b^* mice show a decrease in their DPOAE response to higher frequency stimuli (≥ 12 kHz, P<0.001, one-way ANOVA) (Fig 2F).

Since Np55 is expressed at the OHC stereocilia, we investigated the ability of *Nptn^tm1b^* mice to elicit mechanoelectrical transducer (MET) currents from P7 and P8 OHCs (Fig 3A-H) by displacing their hair bundles in the excitatory and inhibitory direction using a piezo-driven fluid-jet [2, 16]. This age range was selected because it is a time when MET recordings are very reliable and the MET current has reached a mature-like size [17]. At hyperpolarized membrane potentials, the displacement of the hair bundle in the excitatory direction (i.e. towards the taller stereocilia) of P8 OHCs elicited an inward MET current from both *Nptn^+/tm1b^* (Fig 3A) and *Nptn^tm1b/tm1b^* mice (Fig 3B). The maximal MET current in *Nptn^tm1b/tm1b^* P8 OHCs (−1219 ± 138 pA *n* = 9) was significantly reduced compared to that recorded in *Nptn^+/tm1b^* P8 OHCs (−1622 ± 194 pA at −124 mV, *n* = 14, *P* < 0.0001, *t*-test, Fig 3C,E). However, just a day earlier, at P7, the maximum MET current was not significantly different between the genotypes (*P* = 0.7743, one-way ANOVA, Fig 3D, E). Because the MET current reverses near 0 mV, it becomes outward when excitatory bundle displacements are applied during voltage steps positive to its reversal potential (Fig 3A-D). At P8, but not at P7, the maximum MET current was significantly reduced in *Nptn^tm1b/tm1b^* compared to that recorded in *Nptn^+/tm1b^* (P8: *P* < 0.0001, *t*-test; P7: *P* = 0.9537, one-way ANOVA, Fig 3F).

**Fig 3.**
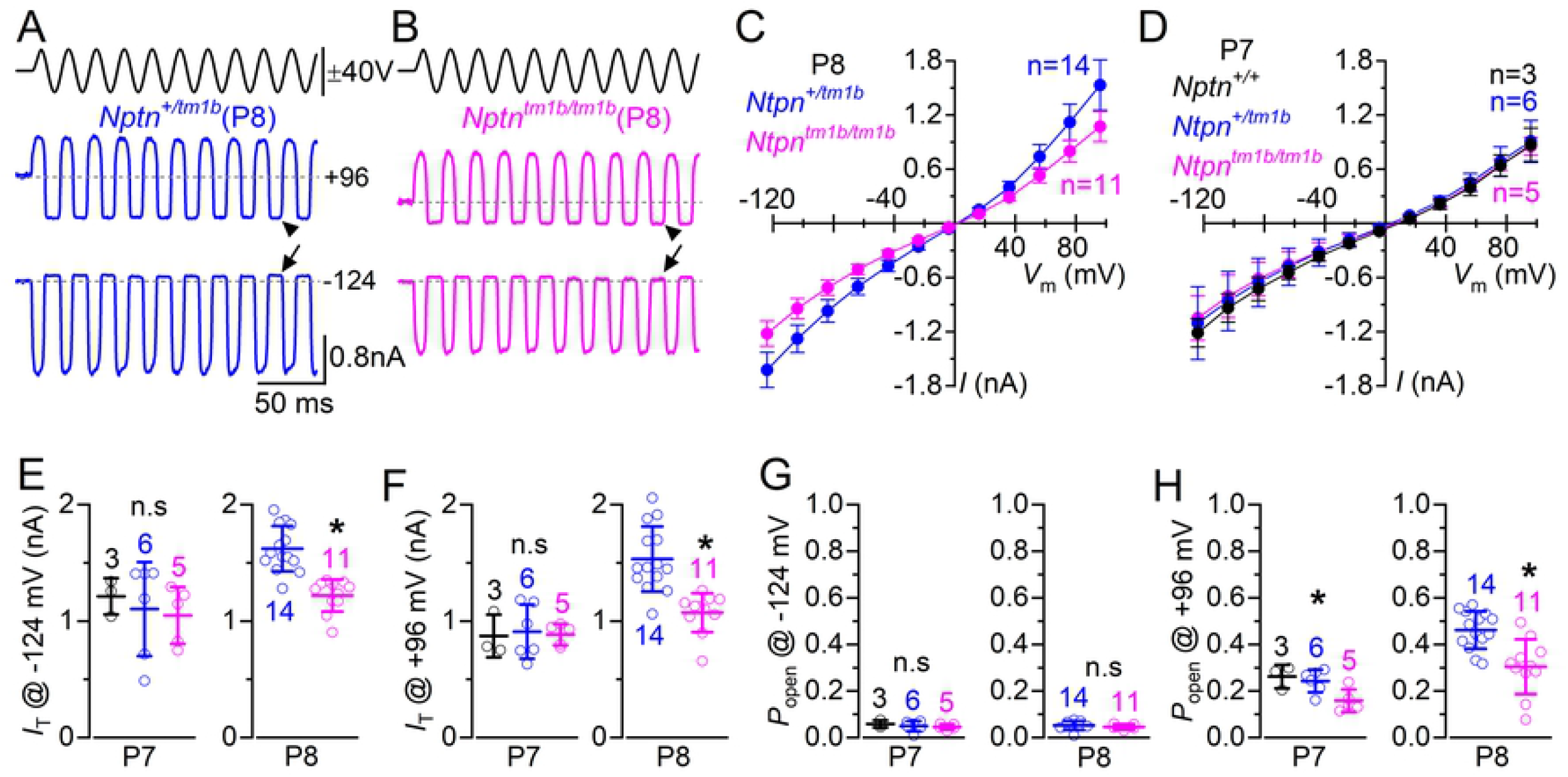
Mechanoelectrical transducer current is affected in *Nptn^tm1b^* mice. (A, B) Saturating MET currents in apical OHCs from a heterozygous *Nptn^+/tm1b^* (A, P8) and a homozygous knockout *Nptn^tm1b/tm1b^* (B, P8) mouse in response to 50 Hz sinusoidal force stimuli to the hair bundles at membrane potentials of −124 mV. Driver voltage (DV) stimuli to the fluid-jet are shown above the traces (excitatory stimuli: positive deflections of the DV). The arrows and arrowheads indicate the closure of the transducer channel in response to inhibitory bundle stimuli at −124 mV and +96 mV, respectively. (C,D) Average peak to peak MET current-voltage curves recorded from P8 (C) and P7 (D) OHCs of mice from the different genotypes indicated in the panels. Currents were obtained by stepping the OHC membrane potential from −124 mV to +96 mV in 20 mV increments while mechanically stimulating their hair bundles. (E, F) Average maximal MET current recorded at −124 mV (E) and +96 mV (F) from OHCs at P7 and P8 of *Nptn^+/tm1b^* and littermate *Nptn^tm1b/tm1b^* mice. Single-data points are shown as open symbols. (G, H) Resting open probability (*P_o_*) of the MET current at the holding potential of −124 mV (G) and +96 mV (H) measured from P7 and P8 OHCs of both genotypes. The resting current is given by the holding current minus the current present during inhibitory bundle deflection. Data are mean ± SD.

The resting open probability (*P_o_*) of the MET channel, which is defined by the resting MET current flowing through open MET channels in the absence of mechanical stimulation, can be measured from the difference between the holding current and the current present during inhibitory bundle deflection (MET channel closed). At negative membrane potentials (−124 mV), the resting current (Fig. 3A,B, arrows) was present in OHCs and was not significantly different at both P7 (P = 0.6316, one-way ANOVA) and P8 (P = 0.5263, *t*-test) (Fig 3G). At positive potentials (+96 mV), the MET channels showed a larger resting transducer current (Fig 3A, B, arrowheads), which is due to an increased open probability of the transducer channel, resulting from a reduced driving force for Ca^2+^ influx [2, 17, 18]. This larger resting MET open probability at +96 mV was significantly reduced between *Nptn^+/tm1b^* and *Nptn^tm1b/tm1b^* OHCs at P8 (P = 0.0006, *t*-test) and between both *Nptn^+/tm1b^* and *Nptn^tm1b/tm1b^* and wild type OHCs at P7 (P = 0.0201, one-way ANOVA) (Fig 3H). These data imply that the reduced expression or absence of NEUROPLASTIN caused a reduction in the size and resting MET current in OHCs starting from about P8.

We then investigated whether the absence of NEUROPLASTIN caused any additional phenotype in the basolateral membrane of the OHCs since it was also expressed in their cell body. Immature OHCs express a delayed rectifier K^+^ current, named *I_K_* [19], which was present in all mice investigated, irrespective of genotype (Fig 4A-D). The size of the steady-state K^+^ current, measured at 0 mV, did not change significantly between *Nptn^+/+^* (3.16 ± 1.0 nA, n = 5), *Nptn^+/tm1b^* (3.50 ± 0.9 nA, n = 7) and *Nptn^tm1b/tm1b^* (2.95 ± 0.5 nA, n = 6) OHCs (P = 0.5051, one-way ANOVA). At the onset of function (~P8: [19]), OHCs start to down-regulate the immature *I_K_* and instead up-regulate their adult-like K^+^ current, named *I_K,n_,* which is carried by KCNQ4 channels. The size and time-course of *I_K,n_* in the OHCs from heterozygous mice (Fig 4E, F) were indistinguishable from that previously reported in wild type mice [19, 20]. However, in age-matched *Nptn^tm1b/tm1b^* mice, while *I_K_* was no longer present in mature OHCs, the size of the *I_K,n_* current was significantly reduced compared to that measured in *Nptn^+/tm1b^* mice (P < 0.0001, Fig 4E-G). These findings indicate that although OHCs from *Nptn^tm1b/tm1b^* mice are initially able to develop toward mature mechano-sensory receptors, the level of expression of their characteristic basolateral K^+^ current *I_K,n_* remained very low, which indicates that OHCs were unable to reach full maturity. The abnormal MET current and reduced *I_K,n_* in OHCs from *Nptn^tm1b/tm1b^* mice would largely impact their ability to generate physiological receptor potentials, thus explaining the loss of DPOAEs (Fig 1F).

**Fig 4.**
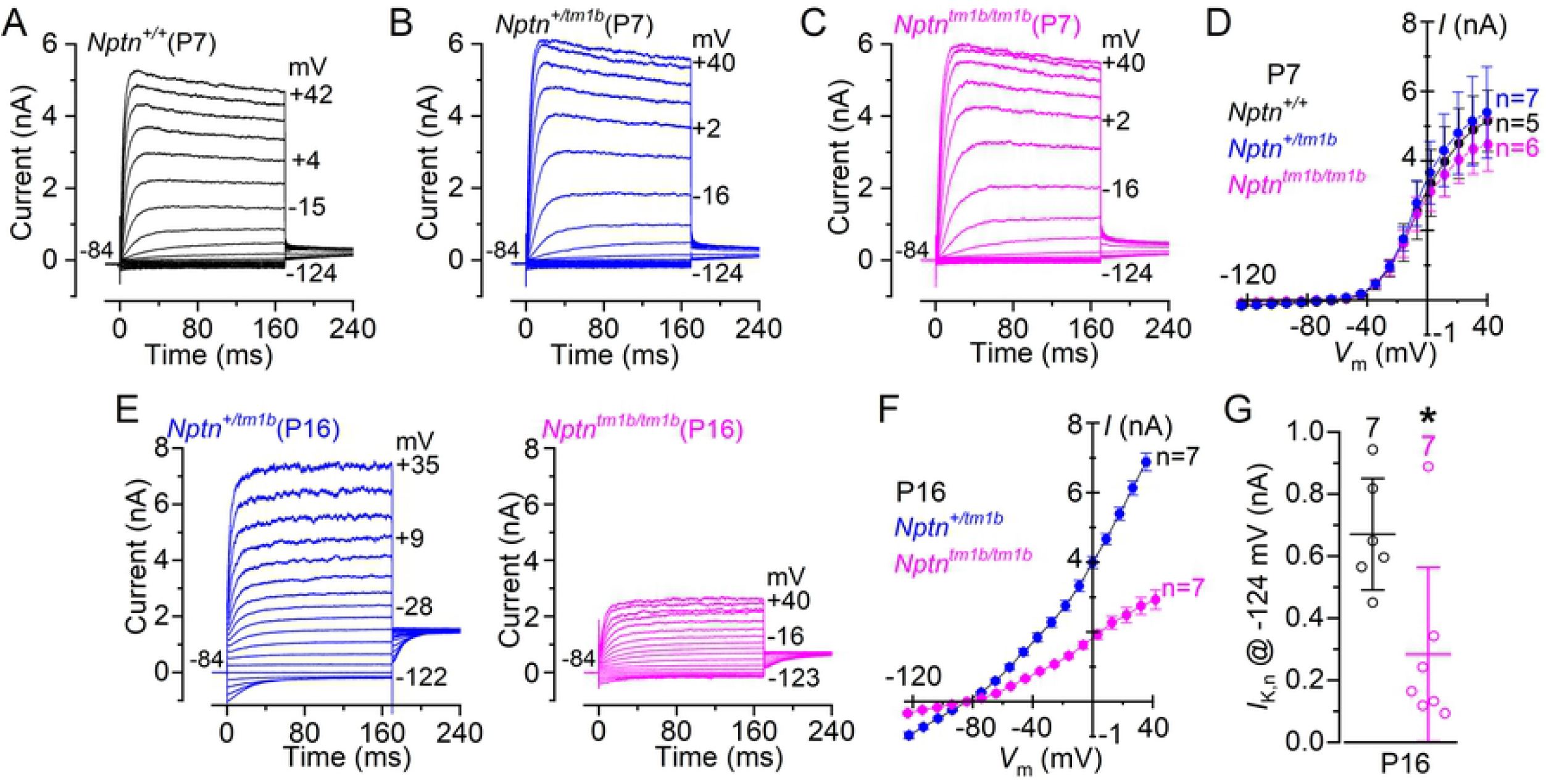
The development of OHCs is disrupted in *Nptn^tm1b^* mice. (A-C) Current responses from OHCs of control (A: *Nptn^+/+^*) heterozygous (B: *Nptn^+/tm1b^*) and homozygous (C: *Nptn^tm1b/tm1b^*) mice at P7, which is during pre-hearing ages. Current recordings were elicited by using depolarising and hyperpolarising voltage steps (10 mV increments) from the holding potential of −84 mV to the various test potentials shown by some of the traces. (D) Steady-state current-voltage curves obtained from P7 OHCs from the three genotypes. (E) Current responses from mature OHCs of *Nptn^+/tm1b^* (left) and *Nptn^tm1b/tm1b^* (right) mice at P16. Note that the time-course of the currents are comparable between the two genotypes, although the current is largely reduced in the *Nptn^tm1b/tm1b^* mouse. (F) Steady-state current-voltage curves obtained from P16 OHCs from the two genotypes. (G, H) Size of *I_K,n_* measured as the difference between the peak and steady-state of the deactivating inward current at −124 mV [19]. The number of IHCs recorded are shown above each column. Average data are plotted as mean ± SD.

### NEUROPLASTIN is required for normal IHC development

Considering the substantial hearing loss observed in young-adult *Nptn^tm1b/tm1b^* mice (Fig 2E, F), and the finding that Np65 (and possibly Np55) is expressed in the basolateral region of the IHCs, we investigated whether the absence of NEUROPLASTIN had any role in their normal function (Fig 5A-G). We found that mature IHCs expressed a large outward K^+^ current, which was present in both *Nptn^+/+^* and *Nptn^+/tm1b^* mice (Fig. 5A, D), but was largely reduced in *Nptn^tm1b/tm1b^* mice (Fig 5B, D). A characteristic K^+^ current of mature IHCs is the rapid activating, large conductance Ca^2+^-activated K^+^ current carried by BK channels, named *I_K,f_* [17, 21, 22]. *I_K,f_* was present in IHCs from *Nptn^+/+^* and *Nptn^+/tm1b^* mice, but absent in *Nptn^tm1b/tm1b^* mice (Fig 5A-C). The size of the total outward K^+^ current *I_K_* and the isolated *I_K,f_* was significantly reduced between the IHCs of *Nptn^+/+^* and *Nptn^+/tm1b^* mice and those from *Nptn^tm1b/tm1b^* mice (*I_K_*: P = 0.0099, Fig 5E; *I_K,f_:* P < 0.0001, Fig 5F, one-way ANOVA). Similar to OHCs, mature IHCs also express *I_K,n_* [23, 24]. In IHCs from *Nptn^tm1b/tm1b^* mice, we observed a very small inward current that could be attributed to *I_K,n_* (Fig 4G), which was significantly reduced compared to that from IHCs of *Nptn^+/tm1b^* mice (P = 0.0007, Fig 5G, one-way ANOVA). However, this current could also be a remnant of the immature inward current *I*_K,1_ [25], since IHCs seem to retain all features of immature cells. This indicates that different from OHCs, the development of IHCs from *Nptn^tm1b/tm1b^* mice appears to be stuck at immature stages.

**Fig 5.**
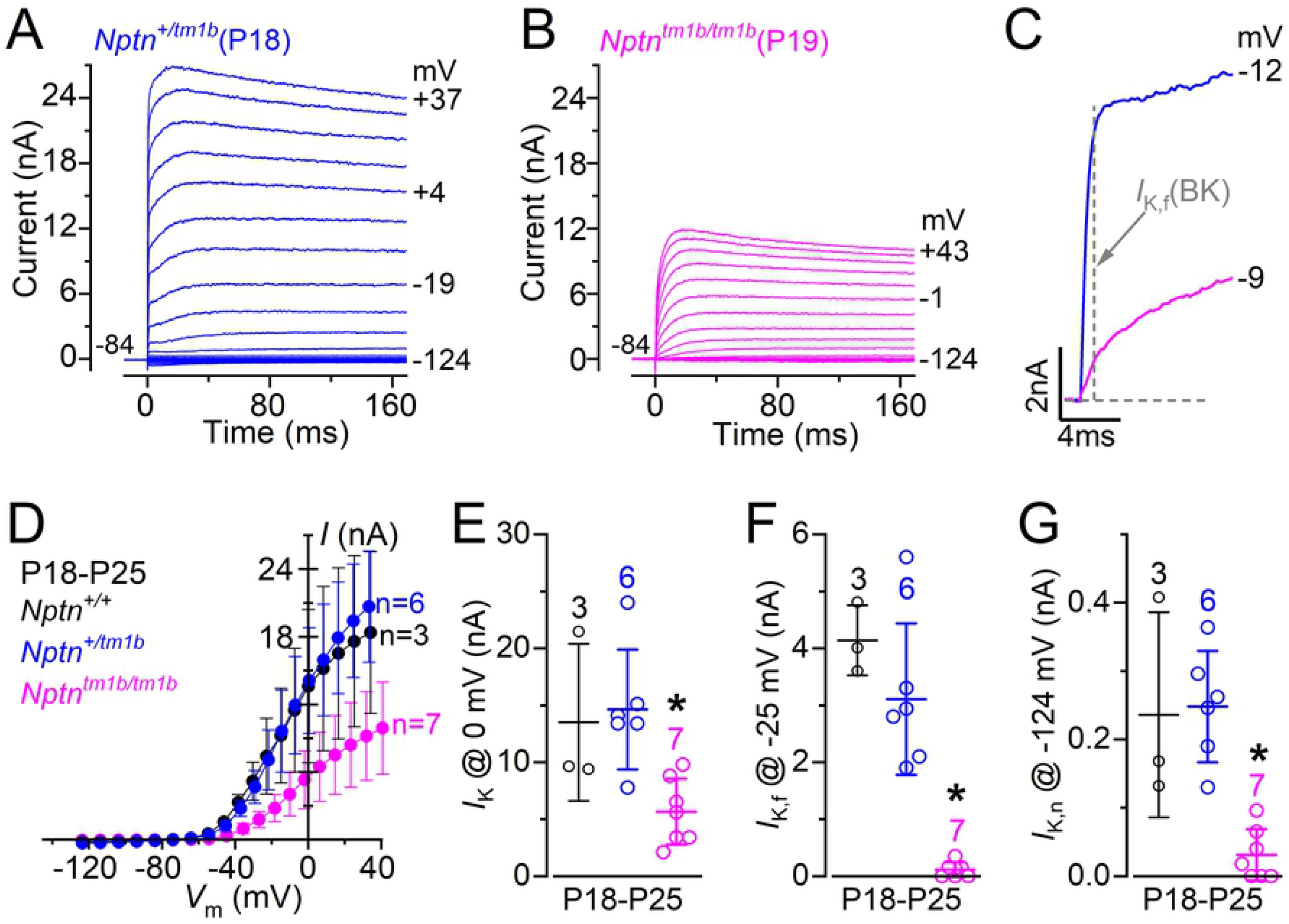
NEUROPLASTIN is required for normal IHC maturation. (A, B) Current responses from IHCs of heterozygous (A: *Nptn^+/tm1b^,* P18) and homozygous (B: *Nptn^tm1b/tm1b^,* P19) mice, which is after hearing onset at P12. Current recordings were elicited by using depolarising and hyperpolarising voltage steps (10 mV increments) from the holding potential of –84 mV to the various test potentials shown by some of the traces. (C) An expanded version of the first 10 ms of the current traces from panels A and B, which emphasises the presence of the rapidly activating *I*_K,f_. (D) Steady-state current-voltage curves obtained from P18-P25 IHCs from the three genotypes. (E-G) Size of the total outward K^+^ current *I*_K_ (E), the isolated *I*_K,f_ (F) and *I*_K,n_ (G). The size of *I*_K,f_ was measured at –25 mV and 1 ms from the onset of the voltage step [22]. The size of *I*_K,n_ was measured as the difference between the peak and steady-state of the deactivating inward current at –124 mV [19]. The number of IHCs recorded are shown above each column. Average data are plotted as mean ± SD.

Previous studies have demonstrated synaptic defects in NEUROPLASTIN deficient mice (*Nptn^pitch^,* [12]). To assess synaptic coupling between IHCs and afferent SGNs in *Nptn^tm1b^* mice, we performed whole-mount immunolabeling of cochleae, using an anti-Ribeye antibody (pre-synaptic ribbon) and an anti-GluR2 antibody (post-synaptic density) (Fig 6A). In the mid-apical region, *Nptn^+/tm1b^* cochleae were found to have similar numbers of matched and unmatched synapses to wild type cochleae, whereas *Nptn^tm1b/tm1b^* cochleae were found to exhibit a 49.0 ± 13.6% reduction in the number of matched pre- and post-synaptic puncta compared to wild type littermates (P < 0.0001, n =3 mice, 30 cells in both genotypes, unpaired *t*-tests with Holm-Sidak correction, Fig 6B). Total counts of Ribeye puncta show that the reduction in matched synapses corresponded to a 44.9 ± 11.5% decrease in the number of IHC ribbons (P < 0.001, n = 3 mice, 30 cells in both genotypes, Fig 6B), suggesting it is loss of the pre-synaptic component that underlies the reduction in matched synapses. To investigate whether a reduction in the number of ribbon synapses may underlie the high-frequency hearing loss exhibited by *Nptn^+/tm1b^* mice (Fig 2E), basal IHC synapses were also assessed. Interestingly, while there was a small decrease in the number of unmatched GluR2 and Ribeye puncta, there was no significant difference in the number of matched synapses when compared with wild type (Fig 6C, D). This suggests that an overt synaptic deficit does not underlie the high-frequency hearing impairment exhibited by the *Nptn^+/tm1b^* mice.

**Fig 6.**
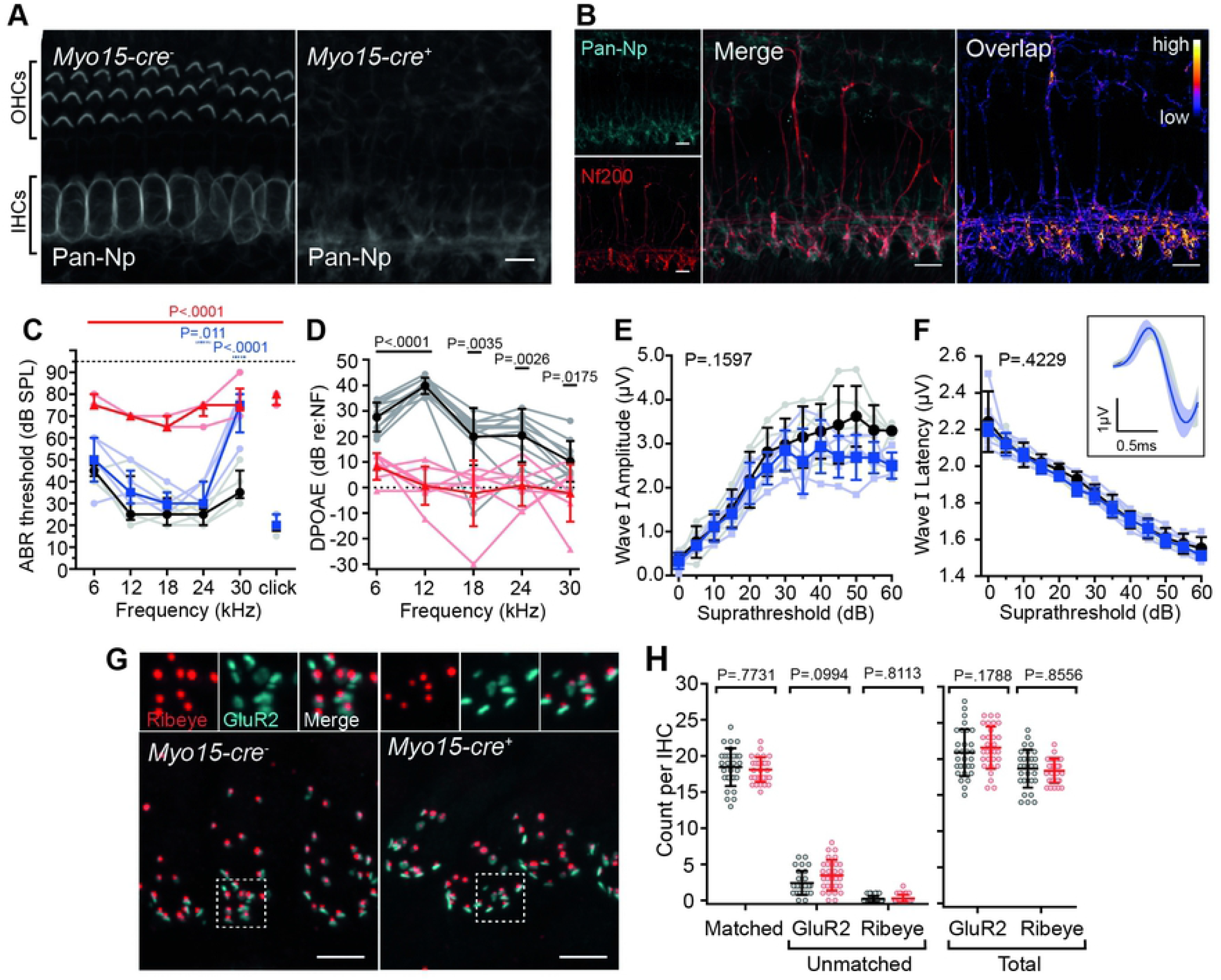
Heterozygous *Nptn^+/tm1b^* mice exhibit reduced ABR wave I amplitudes and increased latencies. (A) Maximum intensity projections of whole-mount mid-apical cochleae from *Nptn^+/+^, Nptn^+/tm1b^,* and *Nptn^tm1b/tm1b^* mice labelled with the pre-synaptic ribbon marker Ribeye (red) and post-synaptic density marker GluR2/3 (cyan). (B) Counts of Ribeye and GluR2/3 puncta were made from ten mid-apical IHCs from *Nptn^+/+^* (black), *Nptn^+/tm1b^* (blue) and *Nptn^tm1b/tm1b^* (red) cochleae (two independent regions per mouse, three mice per genotype). Ribeye and GluR2/3 puncta were considered matched when directly juxtaposed to one another. A larger number of unmatched GluR2/3 puncta and fewer Ribeye puncta were observed in *Nptn^tm1b/tm1b^* cochleae. (C) Maximum intensity projections of whole-mount basal cochleae from *Nptn^+/+^* and *Nptn^+/tm1b^* mice labelled with the pre-synaptic ribbon marker Ribeye (red) and post-synaptic density marker GluR2/3 (cyan). (D) Counts of Ribeye and GluR2/3 puncta were made from 5-10 IHCs from *Nptn^+/+^* (black) and *Nptn^+/tm1b^* (blue) cochleae (two independent regions per mouse, three mice per genotype). *Nptn^+/tm1b^* mice show a decrease in the number of unmatched GluR2/3 puncta compared with *Nptn^+/+^.* Insets are an enlarged view of a 5 μm x 5 μm region highlighted by the dashed box in the corresponding above image. Scale bar = 5 μm. Data are mean ± SD with individual data points shown. Data are compared against wild type controls using unpaired t-tests with a Holm-Sidak correction for multiple comparisons. (E) Averaged ABR wave I in response to a click stimulus recorded at 30 dB above hearing threshold for *Nptn^+/tm1b^* (blue line, n = 5) and *Nptn^+/+^* (black line, n = 5) mice. Across all intensities, *Nptn^+/tm1b^* mice had reduced amplitudes (F) and increased latencies (G) compared with *Nptn^+/+^* mice. Data are mean ± SD with individual data points shown. Analysed using a two-way ANOVA.

To further interrogate the ABR phenotype exhibited by heterozygous *Nptn^+/tm1b^* mice, we examined their recorded ABR waveforms, which represent the sound stimuli-evoked electrical activity in the auditory nerve and brainstem nuclei. In response to the broadband click stimulus, *Nptn^+/tm1b^* mice did not show an overt hearing threshold increase (Fig 2E). However, they do exhibit deficits in ABR Wave I, which reflects the magnitude and synchronicity of activity in the primary afferent neurons. In particular, *Nptn^+/tm1b^* mice show reduced amplitudes (P < 0.0001, n = 5 for both genotypes, two-way ANOVA) and slower onset to peak (P = 0.0002, n = 5 for both genotypes, two-way ANOVA) compared to *Nptn^+/+^* littermate mice (Fig 6E-G).

### Cochlear hair cell expression of NEUROPLASTIN is essential for hearing

To determine if Neuroplastin expression in cochlear hair cells is essential for hearing, we utilised a floxed *Nptn* allele (*Nptn^fl^*) (Fig 1) crossed with a *Myo15-cre* driver mouse line. This conditional knockout mouse (*Nptn^fl/fl^;Myo15-cre^+^*) allowed us to delete *Nptn* specifically in both OHCs and IHCs from approximately P4 onwards [26], which is several days prior to hair cell functional maturation for OHCs (~P8) and IHCs (~P12). Immunolabeling of cochleae from *Nptn^fl/fl^;Myo15-cre^+^* mice confirmed the absence of NEUROPLASTIN in both OHCs and IHCs. However, labelling remained present adjacent to the basal region of IHCs (Fig 7A). Co-labelling with Neurofilament-200K (NF200) indicates that this labelling is associated with NEUROPLASTIN localisation in post-synaptic SGN afferent fibres (Fig 7B). This is consistent with our wild type cochleae immunolabeling data, as well as available scRNA-seq data suggesting *Nptn* transcripts are present in SGNs and HCs (umgear.org).

**Fig 7.**
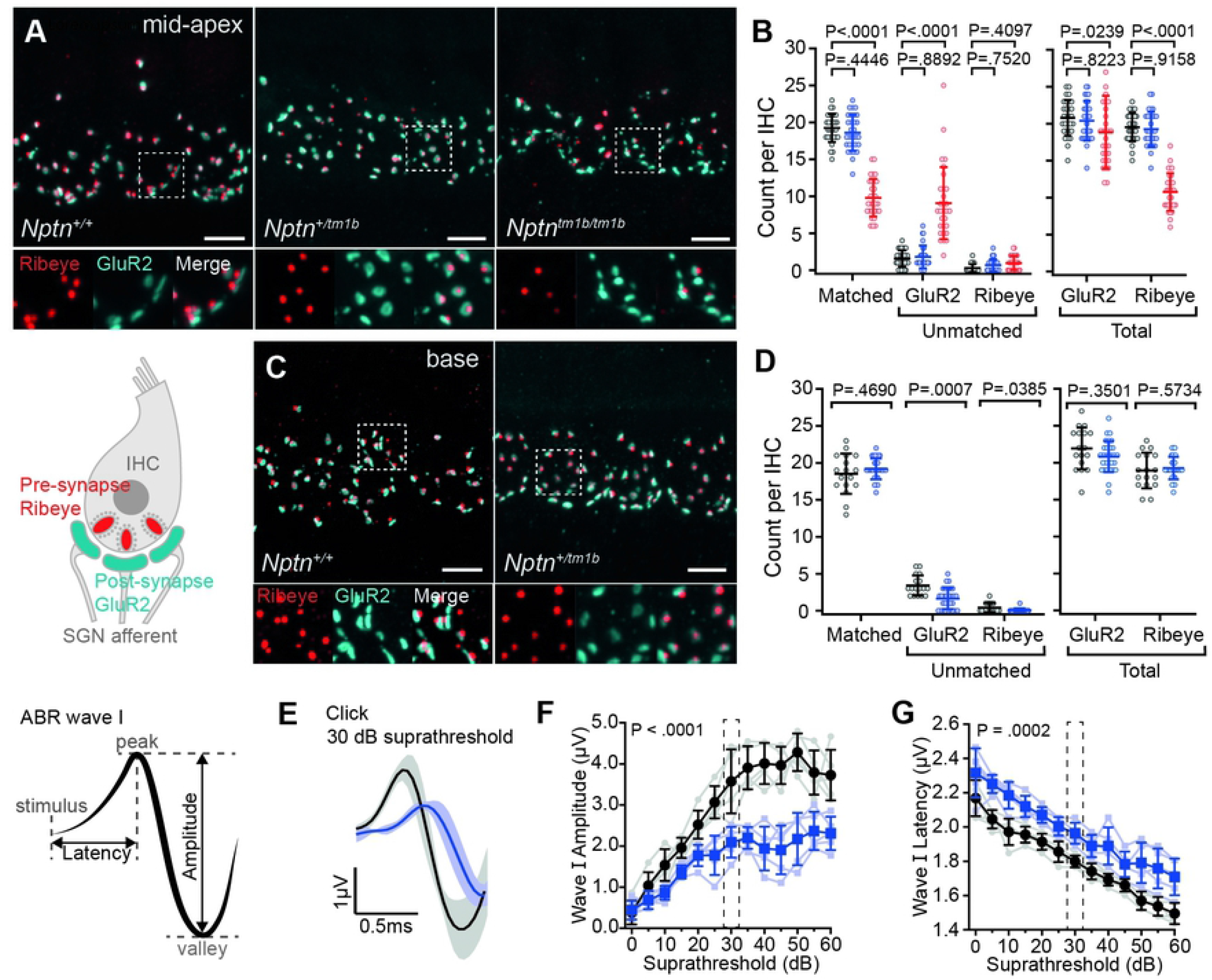
Hair Cell-specific deletion of Neuroplastin elicits a milder auditory phenotype. (A) Maximum intensity projections of whole mount immunohistochemistry of cochleae from 1-month old mice using an anti-pan-Np antibody. Neuroplastin is specifically deleted from both IHCs and OHCs in *Myo15-cre^+^* cochleae. Staining for NEUROPLASTIN is still present around the base of IHCs. (B) Co-labelling with anti-Neurofilament 200K (NF200) in *Nptn^fl/fl^;Myo15-cre^+^* cochleae. Overlap (right) shows regions in which signals for both NEUROPLASTIN and NF200 are detected, with colour indicating summed intensity of signal. Patterns of staining are similar for both antibodies at the base of IHCs suggesting that NEUROPLASTIN labelling is associated with spiral ganglion fibres. Scale bar = 10 μm. (C) ABR threshold measures for *Myo15-cre^+^* mice at 1-month of age shows elevated thresholds in *Nptn^fl/fl^;Myo15-cre^+^* mice (red triangles, n = 5) compared to *Nptn^+/+^;Myo15-cre^+^* mice (black circles, n = 5). Heterozygous *Nptn^+/fl^;Myo15-cre^+^* mice (blue squares, n = 5) also exhibit raised thresholds at ≥ 24 kHz compared with controls. Data are median ± IQR with individual data points shown. One-way ANOVA comparing against wild type using Dunnett’s test for multiple comparisons. (D) Average DPOAE responses of *Nptn^fl/fl^;Myo15-cre^-^* (black circles, n = 10) and *Nptn^fl/fl^;Myo15-cre^+^* (red triangles, n = 7) mice at 1-month of age. Data are mean ± SD with individual data points shown. One-way ANOVA comparing against wild type controls using Dunnett’s test for multiple comparisons. (E-F) Click-evoked ABR wave I measures comparing *Nptn^+/fl^;Myo15-cre^+^* with *Nptn^+/+^;Myo15-cre^+^* mice. Waveforms were not significantly different. Amplitude: P = 0.1597, Latency: P = 0.4229 (two-way ANOVA). Inset shows an average of wave I following click stimulus at 30 dB above threshold. Data are mean ± SD with individual data points shown. (G) Maximum intensity projections of mid-apical cochlear whole-mounts from *Nptn^fl/fl^;Myo15-cre^-^* and *Nptn^fl/fl^;Myo15-cre^+^* mice labelled with the IHC pre-synaptic ribbon marker Ribeye (red) and post-synaptic density marker GluR2/3 (cyan). Insets show an enlarged 5 μm x 5 μm region highlighted by the dashed box in the above image. Scale bar = 5 μm. (H) Counts of Ribeye and GluR2/3 puncta were made from ten adjacent hair cells (two independent regions per mouse, three mice per genotype). Ribeye and GluR2/3 were considered matched when directly juxtaposed to one and other. No significant difference was found in the total numbers of Ribeye or GluR2 puncta, or matched Ribeye/GluR2 puncta. Data are mean ± SD with individual data points shown. Data compared against wild type controls using unpaired t-tests with a Holm-Sidak correction for multiple comparisons.

At four weeks of age, *Nptn^fl/fl^* mice without the cre-driver (*Myo15-cre^-^*) exhibit ABR thresholds similar to those of wild type (*Nptn^+/+^;Myo15-cre^-^*) littermate mice (S1 Fig) demonstrating the floxed allele does not affect hearing. Similarly, wild type mice carrying the *Myo15-Cre (Nptn^+/+^;Myo15-cre^+^*) have normal ABR thresholds (S1 Fig). However, when Neuroplastin is specifically deleted in hair cells (*Nptn^fl/fl^;Myo15-cre^+^*), ABR thresholds are significantly raised (P < 0.0001, one-way ANOVA) to between 65 – 90 dB SPL (Fig 7C), and DPOAEs are reduced across most of the tested frequencies (Fig 7D). Interestingly, the auditory deficit is milder than that exhibited by the global *Neuroplastin* knockout (*Nptn^tm1b/tm1b^*) mice, with no *Nptn^fl/fl^;Myo15-cre^+^* mouse exhibiting an evoked ABR response of >90 dB SPL (NR) at any frequency tested. Similarly, *Nptn^+/fl^;Myo15-cre^+^* mice exhibit a milder hearing loss phenotype than *Nptn^+/tm1b^* mice, with increases in threshold only found at the highest frequency tested (30 kHz, Fig 7C, range: 55 – 80 dB SPL, P < 0.0001, one-way ANOVA). However, in contrast to *Nptn^+/tm1b^* mice, *Nptn^+/fl^;Myo15-cre^+^* mice do not show any differences in the amplitude or latency of ABR wave I following a click stimulus (Fig 7E,F).

Given the continued presence of NEUROPLASTIN at the post-synaptic region, and slightly milder ABR phenotype in *Nptn^fl/fl^;Myo15-cre^+^* mice, we next assessed numbers of Ribeye and GluR2 puncta present in these mice compared with normal hearing *Nptn^fl/fl^;Myo15-cre^-^* mice. No significant difference in the total numbers of Ribeye or GluR2 puncta, or matched Ribeye/GluR2 puncta, were found (Fig 7G, H).

Together, these data show that hair cell expression of *Nptn* is essential for normal hearing function, and suggest that presence of NEUROPLASTIN at the post-synapse, and in the developing HCs up to ~P4, is sufficient to support IHC afferent innervation and ribbon synapse formation.

### Neuroplastin-65 is not required for hearing function

Our finding that *Nptn^tm1b/tm1b^* mice (constitutive null for Np55 and Np65) exhibit ribbon synapse deficits, but that *Nptn^fl/fl^;Myo15-cre^+^* mice (lacking Np55 and Np65 from ~P4) have normal synapses, suggests that NEUROPLASTIN is involved in IHC innervation. Indeed, in a previous paper we propose that Np65 is required for the formation of mature ribbon synapses [12]. Furthermore, our immunolabeling data show that Np65 is present in IHCs (Fig 2D and 8C). However, all of the *Nptn* mutant alleles studied in relation to hearing to date target both Np55 and Np65. To investigate the requirement of Np65 for auditory function, Np65-knockout mice were generated using a CRISPR/Cas9-mediated exon deletion approach, targeting exon 2 of the *Nptn* gene (*Nptn^Δexon2^*) that encodes the Np65-specific Ig1 domain (Fig 1).

**Fig 8.**
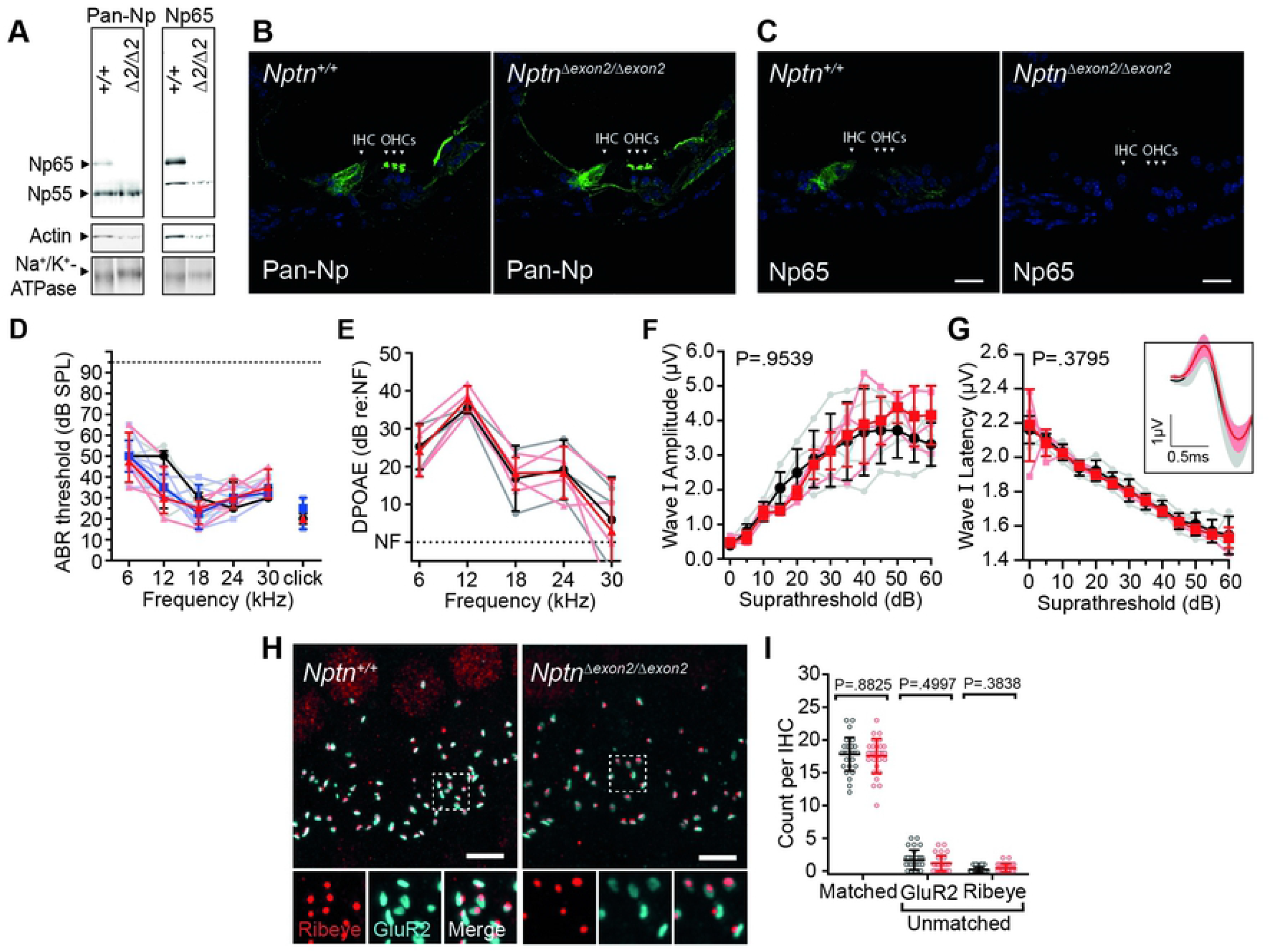
Np65 is not essential for hearing. (A) Western blot of membrane-enriched fractions from *Nptn^+/+^* (+/+) and *Nptn^Δexon2/Δexon2^ (Δ2/Δ2*) cochlear lysates. Lysates were treated with PNGase F to remove N-linked glycans prior to running. The anti-pan-Np antibody detects bands corresponding to Np65 and Np55 in the wild type membrane fraction and the anti-Np65-specific antibody detects a single band corresponding to Np65. No bands were detected in *Nptn^tm1b/tm1b^* lysates when using either antibody. Na^+^/K^+^-ATPase was used as a plasma membrane marker protein. (B) Immunohistochemistry using anti-pan-Np on cochlear cryosections showed labelling of outer hair cell (OHC) stereocilia, the inner hair cells (IHCs), and lateral support cells in both *Nptn^+/+^* and *Nptn^Δexon2/Δexon2^* cochleae. (C) the anti-Np65 specific antibody showed labelling of IHCs in *Nptn^+/+^* cochleae. No signal was observed in *Nptn^Δexon2/Δexon2^* cochleae. Scale = 20 μm. (D) ABR threshold measures of 1-month old mice. *Nptn^Δexon2/Δexon2^* (red triangle, n = 4) and *Nptn^+/Δexon2^* (blue square, n = 6) mice have thresholds similar to *Nptn^+/+^* littermate controls (black circle, n = 5) across all measured frequencies. Data are median ± IQR with individual data points shown. One-way ANOVA comparing against wild type controls with Dunnett’s correction for multiple comparisons. (E) Average DPOAE responses at 1-month of age showing no significant difference between *Nptn^Δexon2/Δexon2^* (n = 4) and littermate controls (*Nptn^+/+^*, n = 3). Data are mean ± SD with individual data points shown. One-way ANOVA comparing against wild type controls using Dunnett’s test for multiple comparisons. (F-G) Click-evoked ABR wave I measures comparing *Nptn^+/+^* (black, n = 5) and *Nptn^Δexon2/Δexon2^* (red, n = 4) mice. Waveforms were not significantly different. Amplitude: P = 0.9539, Latency: P = 0.3795 (two-way ANOVA). Inset shows an average of wave I following click stimulus at 30 dB above threshold. Data are mean ± SD with individual data points shown. (H) Maximum intensity projections of mid-apical cochlear whole-mounts from *Nptn^Δexon2^* mice labelled with the pre-synaptic ribbon marker Ribeye (red) and post-synaptic density marker GluR2/3 (cyan). Insets show an enlarged 5 μm x 5 μm region highlighted by the dashed box in the above image. Scale bar = 5 μm. (I) Ribeye and GluR2/3 puncta count from ten hair cells (two independent regions per mouse, three mice per genotype). Ribeye and GluR2/3 were considered matched when directly juxtaposed to one another. No significant difference was found in *Nptn^Δexon2/Δexon2^* cochleae. Data are mean ± SD with individual data points shown. Data compared against wild type controls using unpaired t-tests with a Holm-Sidak correction for multiple comparisons.

In *Nptn^Δexon2/Δexon2^* mice, the absence of Np65 in cochlear tissue was confirmed by Western blotting, while Np55 remained present (Fig 8A). Immunolabeling of wild type (*Nptn^+/+^*) and *Nptn^Δexon2/Δexon2^* cochleae, utilising the anti-pan-Np antibody, showed no overt changes in signal location or intensity in either IHCs, OHCs, or lateral non-sensory cells (Fig 8B). NEUROPLASTIN expression was also assessed using the anti-Np65-specific antibody, which labelled *Nptn^+/+^* IHCs, but was absent from *Nptn^Δexon2/Δexon2^* IHCs (Fig 8C), thus confirming the successful deletion of Np65 in these mice. ABR recordings showed that at 4-weeks of age *Nptn^Δexon2/Δexon2^* mice have thresholds within the typically normal range (Fig 8D, *Nptn^Δexon2/Δexon2^:* range 15-65 dB SPL, n = 4; *Nptn^+/+^:* range 15-60 dB SPL, n = 5), and their DPOAE responses were also indistinguishable from control littermates (Fig 8E). Furthermore, examination of their ABR waveforms in response to a broadband click stimulus showed no overt deficits in the magnitude and synchronicity of activity in primary afferent neurons (Fig 8F, G). We also assessed IHC ribbon synapses in 3-week old *Nptn^Δexon2/Δexon2^* mice, and found no significant differences in their numbers (Fig 8H, I). To assess whether Np65-null mice exhibit a late-onset progressive hearing impairment, the *Nptn^Δexon2^* allele was first crossed onto a *Cdh23*-repaired C57BL/6N background to circumvent interference from the age-related deafness-causing strain-specific *Cdh23^ahl^* allele [27]. No declines in hearing thresholds were observed up to 6-months of age (Fig 9A-B), and DPOAE responses were comparable over the same time scale (Fig 9C). Together these data suggest that the NEUROPLASTIN isoform Np65 has a redundant role in the establishment and maintenance of mammalian hearing and that Np55 alone is sufficient for auditory function.

**Fig 9.**
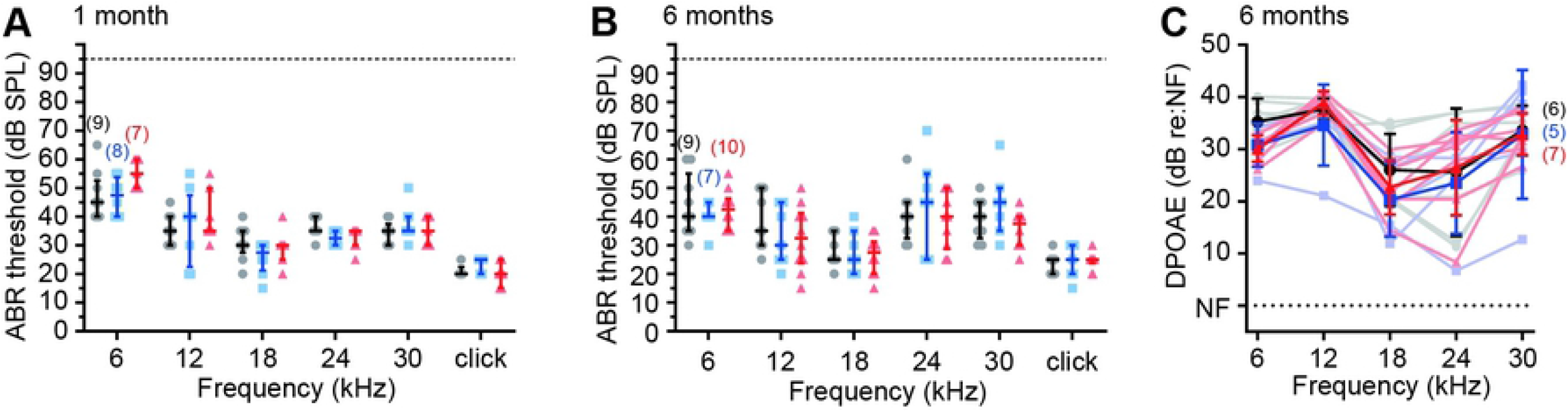
Np65-null mice maintain normal hearing up to 6-months of age. The *Nptn^Δexon2^* allele was crossed onto the *Cdh23*-repaired C57BL/6N background to circumvent interference from the strain-specific *Cdh23 age-related hearing loss* (*Cdh23^ahl^*) allele, and ABR thresholds were tested longitudinally at (A) 1-month and (B) 6-months of age. At each timepoint, the three genotype groups had similar thresholds and were within the expected normal range for the frequencies tested. Data are median ± IQR with individual data points shown. The number of individual mice in each genotype group are shown in brackets. One-way ANOVA comparing against wild type with Dunnett’s correction for multiple comparisons. (C) DPOAE responses of Np65-null (*Nptn^Δexon2/Δexon2^*) mice at 6-months of age are comparable to age-matched wild type littermates. Data are mean ± SD. One-way ANOVA comparing against wild type littermate controls using Dunnett’s test for multiple comparisons.

### OHC-expressed NEUROPLASTIN is required for PMCA2 localisation to stereocilia

Recently, NEUROPLASTIN has been reported to be an obligatory subunit to Plasma Membrane Calcium ATPase (PMCA) proteins, with NEUROPLASTIN being essential for their correct localisation to membranes [28]. Moreover, it was shown *in vitro* that the transmembrane domain of NEUROPLASTIN alone is sufficient to drive the localisation of PMCAs to the membrane [28]. In the auditory system, PMCA2 and PMCA1 are both important for hair cell calcium homeostasis by extruding cytoplasmic Ca^2+^ ions into the extracellular fluid. PMCA2 is expressed abundantly in OHC stereocilia, while PMCA1 is localized to the lateral and basolateral membrane of IHCs [29]. Using immunolabeling of wholemount wild type cochleae, we also find PMCA2 and PMCA1 labelling at these locations, as well as in the lateral non-sensory cells (Fig 10A). However, in *Nptn^tm1b/tm1b^* mutants, PMCA immunoreactivity in IHCs, OHCs and non-sensory cells was markedly reduced, although not completely absent (Fig 10B), which corresponds to a reduction of both PMCA2 and PMCA1 (Fig 10A). However, there was no evidence of reduced PMCA immunoreactivity in *Nptn^Δexon2/Δexon2^* cochleae (Fig 10C).

**Fig 10.**
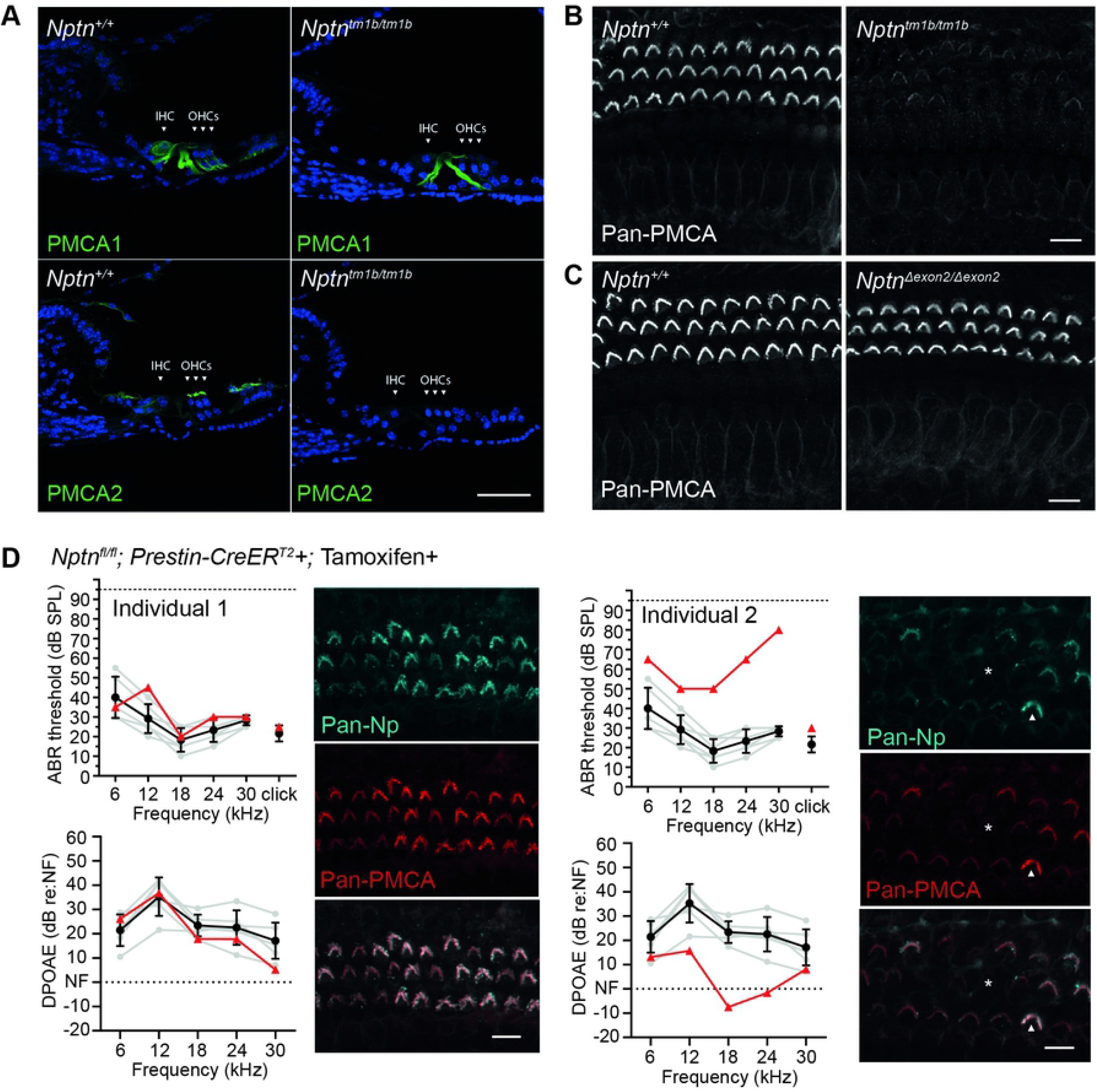
Hearing impairment is inversely correlated with Nptn-dependent localisation of PMCA in outer hair cell stereocilia. (A) Immunolabelling of cochlear cryosections using anti-PMCA1 and anti-PMCA2 antibodies. In wild type tissue, PMCA1 is predominantly localised to the membrane of IHCs and OHCs, whereas PMCA2 is localised to OHC stereocilia and lateral supporting cells. Labelling of the PMCAs at these locations was substantially reduced in *Nptn^tm1b/tm1b^* cochleae. Scale = 20 μm. (B) The anti-pan-PMCA antibody detects PMCA1-4. Maximum intensity projections of whole-mount cochlear immunolabelling using the anti-pan-PMCA antibody showed labelling of OHC stereocilia and membrane of IHCs in *Nptn^+/+^* mice, which were substantially reduced, but not absent, in *Nptn^tm1b/tm1b^* cochleae. (C) Using the anti-pan-PMCA antibody, no labelling differences were seen in *Nptn^Δexon2/Δexon2^* cochleae compared to wild-type. Scale = 10 μm. (D) ABR threshold and DPOAE response measures from two *Nptn^fl/fl^;Prestin-CreER^T2^+* mice following tamoxifen-induced cre-mediated recombination, with corresponding anti-pan-Np and anti-pan-PMCA immunolabelled mid-apical cochlear whole mounts (maximum intensity projections). Data from the individual mice are shown (red triangles) compared against wild type controls (black circles, mean ± SD). Due to varying levels of induction, the auditory phenotype exhibited was variable with some mice showing no significant phenotype (e.g. individual 1), while others showed increased ABR thresholds and a decreased DPOAE response (e.g. individual 2). The auditory phenotype was directly correlated to the level of *Nptn* recombination as demonstrated by immunolabelling using the anti-pan-Np antibody. Bundles that were strongly stained for NEUROPLASTIN also had strong labelling for anti-pan-PMCA antibody (arrowhead), whereas absent NEUROPLASTIN was correlated with a weak PMCA signal (*).

To further examine the requirement of NEUROPLASTIN for the localisation of PMCA2 in OHCs, we generated *Nptn^fl/fl^*;Prestin-CreER^T2^ mice, and induced cre-mediated recombination specifically in OHCs through tamoxifen delivery at 4-weeks of age. When tested by ABR and DPOAE 10-14 days following tamoxifen administration, these mice were found to have a highly variable phenotype with the degree of auditory deficit being directly correlated with the number of NEUROPLASTIN-negative OHCs. Auditory function was not affected by the presence of the Prestin-CreER^T2^ allele or the delivery of tamoxifen alone (S1 Fig). Furthermore, in *Nptn^fl/fl^*;Prestin-CreER^T2^ mice PMCA localisation was directly correlated with NEUROPLASTIN expression, with NEUROPLASTIN positive OHCs showing stronger labelling for PMCAs in stereocilia than adjacent OHC bundles lacking NEUROPLASTIN expression (Fig 10D). These data are consistent with a recently published study by Lin *et al* who show reduced cochlear expression of PMCAs in a different *Neuroplastin* null mouse mutant (*Nptn^tm1.2Mtg^,* [15]).

Our data confirm that NEUROPLASTIN is required for the correct localisation of PMCA proteins in the organ of Corti, and also shows that continued expression of *Neuroplastin* in the OHCs of adult mice is required to maintain hearing function.

### Neuroplastin genetically interacts with *Cdh23^ahl^* allele in C57BL/6 mice

To date, there have only been reports of recessive auditory phenotypes in *Neuroplastin* mutant mice [12, 13]. However, here we observe significant high-frequency hearing loss in both *Nptn^+/tm1b^* and *Nptn^+/fl^;Myo15-cre^+^* mice (Fig 2E and 7C), with a corresponding reduction in DPOAEs seen when testing *Nptn^+/tm1b^* mice (Fig 2F). Given we find that NEUROPLASTIN is required for localisation of PMCA2 to OHC stereocilia (Fig 10), and previous studies show a genetic interaction between *Atp2b2* (encodes PMCA2) mutations and the hypomorphic *Cdh23^ahl^* allele present in the C57BL/6 background [30], we tested whether the *Cdh23^ahl^* allele is a genetic modifier of *Neuroplastin.* To enable this, we crossed the *Nptn^tm1b^* allele onto a C57BL/6N background in which the *Cdh23^ahl^* allele has been corrected (*Cdh23^753A>G^)[27]. Nptn^tm1b/tm1b^* mice on the repaired C57BL/6N (*Cdh23^753A>G/753A>G^*) background exhibit a hearing phenotype comparable to *Nptn^tm1b/tm1b^* mutants on the standard C57BL/6N (*Cdh23^ahl/ahl^*) background when tested by ABR at 4-weeks, and DPOAE at 5-6 weeks (Fig 11A, B). In contrast, *Nptn^+/tm1b^* mice on the repaired C57BL/6N background did not exhibit high-frequency hearing loss, having thresholds similar to wild type control littermates (Fig 11A, B). Directly comparing hearing thresholds of *Nptn^+/tm1b^* mice on the standard C57BL/6N background to age-matched *Nptn^+/tm1b^* mice on the repaired C57BL/6N background, shows that on the standard background, *Nptn^+/tm1b^* mice have significantly elevated thresholds at 18, 24, and 30 kHz (P < 0.0001, one-way ANOVA). Furthermore, while *Nptn^+/tm1b^* mice maintained on a standard C57BL/6N (*Cdh23^ahl/ahl^*) background show significant deficits in click-evoked ABR wave I compared to their wild type littermates (Fig 6E-G), *Nptn^+/tm1b^* mice maintained on the repaired C57BL/6N background do not show any wave I amplitude or latency differences compared to their wild type littermates (Fig 11C, D).

**Fig 11.**
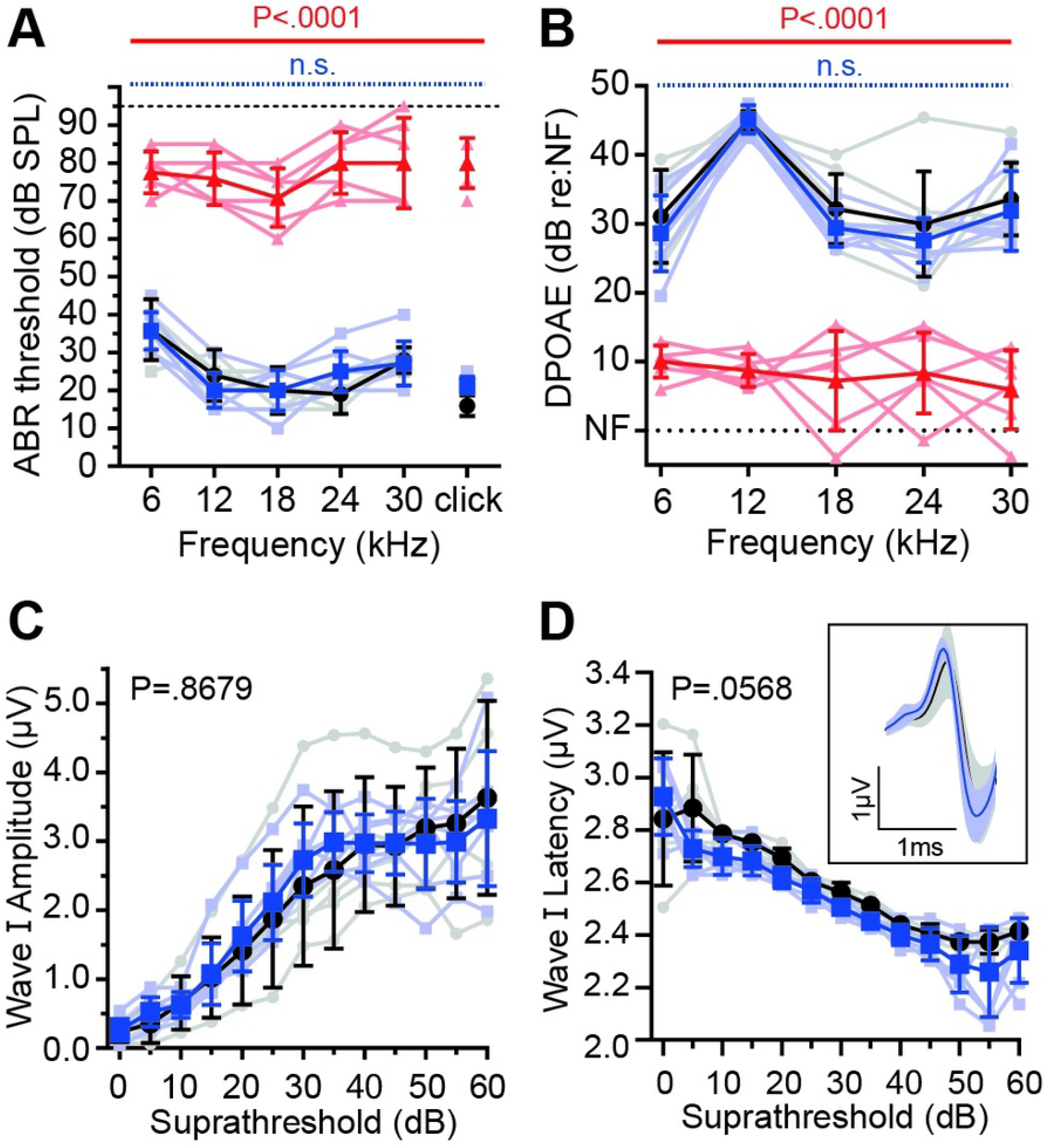
Neuroplastin genetically interacts with Cadherin 23. Phenotypic measures of *Nptn^tm1b^* mice maintained on a *Cdh23*-repaired C57BL/6N (*Cdh23^753A>G^*) background. (A) ABR threshold measurements recorded from 4-week old mice. Similar to *Nptn^tm1b/tm1b^* mice on the standard C57BL/6N genetic background, *Nptn^tm1b/tm1b^* mice on the repaired background (red triangles, n = 6) have profoundly elevated thresholds compared with littermate *Nptn^+/+^* mice (black circles, n = 5). However, on the repaired background *Nptn^+/tm1b^* (blue squares, n = 7) have thresholds indistinguishable from control mice, even at the higher frequencies. Data are median ± IQR with individual data points shown. One-way ANOVA comparing against wild type controls using Dunnett’s test for multiple comparisons. (B) DPOAE measurements recorded at 5-6-weeks of age. Response amplitudes were significantly reduced in *Nptn^tm1b/tm1b^* mice (red triangles, n = 5), whereas *Nptn^+/tm1b^* mice (blue squares, n = 5) have amplitudes comparable to that of control mice (*Nptn^+/+^,* black circles, n = 5). Data are mean ± SD with individual data points shown. One-way ANOVA comparing against wild type controls using Dunnett’s test for multiple comparisons. (C,D) Click-evoked ABR wave I comparing *Nptn^+/+^* (black, n = 5) and *Nptn^+/tm1b^* (blue, n = 7) mice. Waveforms were not significantly different. Amplitude: P = 0.8679, Latency: P = 0.0568 (two-way ANOVA). Inset shows an average of wave I following a click stimulus at 30 dB above threshold. Data are mean ± SD with individual data points shown.

Our data reveal a genetic interaction between *Neuroplastin* and *Cadherin 23* (Otocadherin), highlighting the importance of understanding and reporting the genetic background of mutant mouse models to allow interpretation of phenotypic expressivity and reproducibility of data, respectively.

## Discussion

The expression, localisation and function of Neuroplastin in the mammalian cochlea is a subject of conflicting reports. Here, we show that both Np65 and Np55 are present in the murine cochlea and that the two isoforms exhibit different patterns of cellular expression. Using a pan-Np antibody, we find NEUROPLASTIN to be present in OHC stereocilia, IHC basolateral membrane, spiral ganglia, and also in some lateral non-sensory cells, which is in broad agreement with previous studies [12, 13, 15]. To discern the contribution to this pan-Np labelling arising from Np65, we generated a Np65-specific knockout (*Nptn*^Δ*exon*2^) mouse model. Utilising tissues derived from these mice we were able to confirm the specificity of a Np65-specific antibody, and infer localisation of Np65 at the basolateral region of IHCs and some lateral non-sensory cells within wild type tissues. Importantly, no Np65 labelling was observed in OHC stereocilia, supporting the findings of Zeng, et al. [13], and showing that Np55 is the sole NEUROPLASTIN isoform found in these structures. Although a previous report had suggested that transcripts for Np65 are present in SGNs [13], we found no evidence of the “neuron-specific” Np65 in SGNs by immunolabeling (Figs 2 and 8). We had previously hypothesised that the presence of Np65 at the basolateral membrane of IHCs may act to establish the afferent synapses through *trans*-homophilic dimerisation across the synaptic cleft, mediated via the Np65-specific Ig1 domain. However, when Np65 is specifically deleted in *Nptn^Δexon2^* mice, they showed no observable auditory phenotype, and more importantly, no overt changes to the ribbon synapse count or structure (Fig 8). While we show Np65 is present in wild type IHCs, and Np55 is present in IHCs of Np65 knockout (*Nptn^Δexon2/Δexon2^*) mice, we were not able to determine if Np55 is natively expressed in wild type IHCs, or if is only expressed in place of Np65 in the Np65 knockout. Regardless, the lack of an auditory phenotype in homozygous *Nptn^Δexon2^* mice leads us to conclude that Np65 is functionally redundant in the mammalian cochlea. Thus, we can conclude that Np65-Np65 *trans*-homophilic dimerisation is not a mechanism by which afferent contacts are established and maintained at the IHC ribbon synapse. This does not exclude other mechanisms by which NEUROPLASTIN could promote neurite outgrowth, such as the activation of FGFR in the early development of synapses, which Np55 is capable of through cis-heterophilic binding [7]. Unfortunately, given the structure of the Neuroplastin gene, we are unable to generate a Np55-specific knockout and therefore are not able to determine if Np65 alone would be able support auditory function.

### NEUROPLASTIN expression in hair cells is essential for hearing

To investigate the requirement of NEUROPLASTIN in auditory function, we utilised the *Nptn^tm1a(EUCOMM)Hmgu^* mice that were generated as part of the IMPC programme (https://www.mousephenotype.org/). This is a knockout-first allele that enables the generation of a *Nptn^tm1b(EUCOMM)Hmgu^* knockout allele through cre-mediated recombination (herein referred to as *Nptn^tm1b^),* or a *Nptn^tm1c(EUCOMM)Hmgu^* conditional allele through flp-mediated recombination (herein referred to as *Nptn^fl^*) (Fig 1). Preliminary auditory phenotyping of four homozygous *Nptn^tm1b(EUCOMM)Hmgu^* mice, at 14-weeks of age, identified this as a hearing allele [11]. Using a larger cohort of mice we have validated and confirmed that when homozygous the *Nptn^tm1b^* allele results in an early-onset profound hearing loss, with mice exhibiting thresholds similar to those reported for three ENU-induced mutants: *pitch* (C315S), Y219X, and *audio-1* (I122N), affecting exons 6, 4, and 3, respectively [12, 13]. Furthermore, additional testing of the *Nptn^tm1b/tm1b^* mice showed that DPOAE responses were significantly reduced (Fig 2), indicating OHC dysfunction as also reported for the *Nptn^audio-1^* model [13], and an IHC synaptic deficit comprising of unmatched pre- and post-synaptic markers (Fig 6), as also reported for the *Nptn^pitch^* model [12]. Interestingly, unlike previously reported *Nptn* mutant mouse models, we find an auditory phenotype in heterozygous *Nptn^tm1b/+^* mutant mice. While the phenotype is less severe than that exhibited by homozygous *Nptn^tm1b/tm1b^* mice, it does result in significantly elevated thresholds for higher frequency stimuli (≥18 kHz) and correspondingly reduced DPOAE responses (≥12 kHz). A deficit was also apparent in click-evoked ABR wave I amplitudes and latencies, despite no elevation of thresholds with this stimulus or IHC synaptic deficits. Furthermore, utilising the floxed mice in combination with a hair cell-specific cre-driver line, we show that deletion of *Nptn* in hair cells leads to elevated hearing thresholds in *Nptn^fl/fl^;Myo15-cre^+^* and *Nptn^fl/+^;Myo15-cre^+^* mice, but that their hearing loss is notably milder than that exhibited by aged-matched *Nptn^tm1b/tm1b^* and *Nptn^tm1b/+^* mice, respectively (Fig 7). Moreover, also different to *Nptn^tm1b/tm1b^* mice, *Nptn^fl/fl^;Myo15-cre^+^* mice do not exhibit an observable synaptic deficit, despite having increased auditory thresholds. Taken together, our data show that the expression of Neuroplastin in hair cells is essential for hearing, and suggest that expression of NEUROPLASTIN in the SGNs of hair cell-specific *Nptn* knockout mice is sufficient for the establishment and maintenance of IHC ribbon synapses. Interestingly, in both macaque and guinea pig, the estimated contribution of OHCs to hearing thresholds as a cochlear amplifier is around 50 dB [31]. Here, threshold elevations in *Nptn^tm1b/tm1b^* mice are as high as 69 ± 6 dB (Click), while in the mutant models which do not have changes to synapse structure (i.e. *Nptn^+/tm1b^* and *Nptn^fl/fl^;Myo15-cre^+^*) maximum threshold shifts are 50 ± 10 dB (30 kHz) and 56 ± 4 dB (click), respectively. This suggests that both a reduction in OHC function and disruption to IHC synapses are likely to contribute to the auditory phenotype seen in the NEUROPLASTIN-null *Nptn^tm1b/tm1b^* mice.

### NEUROPLASTIN is essential for hair cell mechanotransduction, maturation and ion homeostasis

Despite localisation of NEUROPLASTIN in OHC stereocilia, OHC MET currents have previously been reported as unaffected in Neuroplastin loss-of-function mutants [12, 13]. However, these were recorded from immature OHCs (≤P7). Here, we find that OHCs of P8 *Nptn^tm1b/tm1b^* mice have significantly reduced maximal MET currents and MET channel open probability at depolarised potentials, which is not evident at P7 (Fig 3). When stepping the OHC membrane potential to positive values, which is near the Ca^2+^ equilibrium potential, the drive for Ca^2+^ entry into the MET channels is strongly reduced. This reduced Ca^2+^ influx leads to the MET channels’ increase open probability, which is a manifestation of the ability of Ca^2+^ to drive adaptation and thus closing the MET channel, as demonstrated in hair cells from lower vertebrates [32–34] and mouse cochleae [2]. Interestingly, we also show that *Nptn^tm1b/tm1b^* mice have significantly reduced membrane-localisation of plasma membrane Ca^2+^ ATPase (PMCA) proteins across the organ of Corti (Fig. 10), which has been reported in other tissue types [35, 36] and more recently in cochleae using a different NEUROPLASTIN-null mouse, *Nptn^tm1.2Mtg^* [15]. Since the expression of the Ca^2+^ pump PMCA is largely reduced at the stereocilia of OHCs from *Nptn^tm1b/tm1b^* mice, this would most likely cause Ca^2+^ to accumulate intracellularly near the MET channel during repetitive bundle stimulation, leading to its adaptation and a reduced open probability compared to wild type OHCs. Although MET channel recordings from PMCA knockout mice is limited, some evidence for a similar reduction in the MET channel open probability has previously been obtained from newborn mouse OHCs lacking the Ca^2+^ pump [37]. In addition to the defects in the MET apparatus, we find that both OHCs and IHCs of *Nptn^tm1b/tm1b^* mice fail to develop a fully mature basolateral current profile (Figs 4 and 5), which is in line with previous work investigating other mutations that affect mechanotransduction, such as mutations in *Eps8* [38], TMC1 [39], *Pcdh15, Harmonin* [40] and *Clrn2* [41]. IHCs from *Nptn^tm1b/tm1b^* mice retain a full pre-hearing basolateral current profile, which could be an indirect consequence of impaired mechanotransduction, as seen in P8 OHCs. This is because depolarising MET currents are critical for driving spontaneous action potential activity in IHCs during pre-hearing stages [42], a key physiological aspect required for IHC maturation [40]. Different from IHCs, OHCs from *Nptn^tm1b/tm1b^* mice are able to mature albeit expressing a reduced K^+^ current. The possible mechanism linking the observed defects in the MET current and the reduced expression of *I*_K,n_ is less clear. Knockout mice for the TMC1 channel, Eps8 and Clrn2 have abnormal OHC MET current from about P8 [43], P8 [38] and P6 [41], respectively, but have opposite effects on the basolateral current profile in OHCs (TMC1: no *I*_K,n_ expression; Eps8 and Clrn2: normal *I*_K,n_ expression). To further investigate the requirement of NEUROPLASTIN for the localisation of PMCA proteins, we utilised a tamoxifen-induced OHC-specific Neuroplastin knockout (*Nptn^fl/fl^; Prestin-CreER^T2^*) model and show that continued expression of NEUROPLASTIN in OHCs of adult mice is required for the localisation of PMCAs and the maintenance of normal hearing function (Fig 10). Moreover, reduced expression of NEUROPLASTIN was directly correlated with PMCA2 expression at the membrane, which in turn was correlated with hearing function. However, while membrane localisation of PMCAs was drastically reduced in the absence of NEUROPLASTIN, some PMCA2 could still be detected correctly located at the plasma membrane. This suggests that another protein, which is also able to act as a PMCA subunit, may be partially substituting for the absence NEUROPLASTIN. Interestingly, it is the transmembrane domain of NEUROPLASTIN that is critical for binding to PMCAs [44], and this domain is highly conserved across members of the Basigin group of cell adhesion molecules [45]. As such, this particular role of NEUROPLASTIN could potentially be undertaken by either BASIGIN or EMBIGIN. Notably, *Embigin* was also identified as a hearing loss candidate gene by the IMPC programme [11].

### Neuroplastin genetically interacts with Cadherin 23

Our finding that heterozygous *Nptn* mutant mice exhibit a high-frequency hearing loss has not been reported for other *Nptn* models. Interestingly, this phenotype was completely absent after crossing the *Nptn^tm1b^* allele to a coisogenic background in which the C57BL/6 strain-specific hypomorphic *Cdh23^ahl^* allele has been corrected (*Cdh23^753A>G^*) (Fig 11). The finding that heterozygous *Nptn^tm1b/+^* mutant mice on a ‘corrected’ C57BL/6N background display normal high-frequency hearing thresholds, compared to *Nptn^tm1b/+^* mutant mice on a standard C57BL/6N background, demonstrates a genetic interaction between *Nptn* and *Cdh23.* Mechanistically, this is presumably acting through the previously defined genetic interaction reported between *Atp2b2* (PMCA2) and *Cdh23,* where the hearing loss phenotype exhibited by heterozygous PMCA2 mutant mice (*Atp2b2^+/dfwi5^*) is elevated if one copy of the *Cdh23^ahl^*allele (*+/ahl*) is also present, and further elevated if homozygous (*ahl/ahl*) [30]. Together, these data suggest that having only one wild type copy of *Nptn* (*Nptn^+/tm1b^*) likely causes a reduction of PMCA2 localisation, leading to a hearing loss when compounded by the presence of the *Cdh23^ahl^* allele, and thereby phenocopying what has been reported for *Atp2b2* heterozygous lesions. Given this genetic interaction, it is important to carefully consider the genetic background of *Nptn* mutant mice when investigating the role of NEUROPLASTIN in the auditory system. Moreover, while to date no *NPTN* mutations have been reported in patients with hearing loss, this has focused on searching for homozygous or compound heterozygous *NPTN* lesions.

In humans, mutations in *CDH23* (ENSG00000107736) are associated with early-onset hearing loss (USH1D and DFNB12) [46, 47], and have been implicated in cases of late-onset, progressive hearing loss [48]. Given our data, and that a genetic interaction between *ATP2B2* and *CDH23* has been demonstrated in humans [49], we suggest that patients carrying double *NPTN* and *CDH23* lesions should be further investigated.

In conclusion, our data show that the primary cause of hearing loss in *Nptn*-null mice is due to OHC dysfunction resulting from a reduction of correctly localised PMCA2, and this is exacerbated by the secondary loss of afferent synapses.

## Methods

### Ethical Approval

All animal studies were licensed by the Home Office under the Animals (Scientific Procedures) Act 1986, United Kingdom, and additionally approved by the relevant Institutional Ethical Review Committees (PBF9BD884 to MRB and PCC8E5E93 to WM). Mice were housed under a 12 hour light/12 hour dark cycle and allowed access to food and water ad libitum. All conducted experiments complied with The Journal’s ethics policies.

### Mice

*Nptn^tm1b(EUCOMM)Hmgu^* (hereafter called *Nptn^tm1b^*) were generated as part of the International Mouse Phenotyping Consortium. These mice were produced through cre-mediated conversion of the ‘knockout-first’ tm1a allele, which was achieved by treating IVF derived embryos with a cell-permeable Cre-enzyme. In the converted tm1b allele, exons 5 and 6 of the Neuroplastin gene are deleted, leaving a *lacZ* reporter cassette (Fig 1). Together exons 5 and 6 encode two critical regions, Ig3 and the transmembrane domain, which are present in both *Nptn* splice isoforms; additionally, the *lacZ* cassette contains a splice acceptor that subsumes normal splicing. Thus, no functional protein is expected to be produced from the *Nptn^tm1b^* allele. *Nptn^tm1c(EUCOMM)Hmgu^* (hereafter called *Nptn^fl^*) mice were generated by crossing the tm1a allele to C57BL/6N mice producing Flp recombinase to remove the *lacZ* and *neo* cassette. *Nptn^fl^* mice functionally behave like wild type mice while retaining the LoxP sites flanking exons 5 and 6 (Fig 1B). The generation of conditional knockouts was achieved by crossing *Nptn^fl^* mice to the *Myo15-cre* [26] and *Prestin-creER^T2^* [50] model lines, provided by C. Petit and T. Friedman (Myo15-cre) and J. Zuo (Prestin-creER^T2^).

*Myo15-cre* recombinant mice carry the cre recombinase gene driven by the Myosin-15 gene promoter, which in the cochlea deletes floxed *Nptn* exons 5 and 6 specifically in hair cells from approximately P4 [26]. *Nptn^+/fl^* female mice were crossed with *Nptn^+/fl^;Myo15-cre^+^* males to generate six genotypes (*Nptn^+/+^;Myo15-cre^-^, Nptn^+/fl^;Myo15-cre^-^, Nptn^fl/fl^;Myo15-cre^-^, Nptn^+/+^;Myo15-cre^+^, Nptn^+/fl^;Myo15-cre^+^, Nptn^fl/fl^;Myo15-cre^+^).* For ABR analysis, all six genotypes were tested. To reduce the number of mice generated, subsequent breedings were set up between *Nptn^fl/fl^* and *Nptn^fl/fl^;Myo15-cre^+^* mice to generate *Nptn^fl/fl^* mice with, or without, the cre driver.

Prestin-creERT2 recombinant mice carry the tamoxifen-inducible cre recombinase gene in the Prestin locus following its stop codon, which deletes floxed Nptn exons 5 and 6 specifically in OHCs [50] following delivery of tamoxifen. *Nptn^fl/fl^* females were crossed with *Nptn^fl/fl^*;*Prestin-creER^T2^+* males to generate *Nptn^fl/fl^* mice with or without the cre driver. Recombination was induced by tamoxifen delivered by oral gavage (3 doses delivered over 3 days at 200 mg/kg) at 4 weeks of age, and ABRs and DPOAEs were performed 7, and 14 days following the final tamoxifen dose, respectively.

The isoform-specific mutant, *Nptn^Δexon2^* mutant line was generated by the Molecular and Cellular Biology group at the MRC Harwell Institute. Using a CRISPR–Cas9-mediated deletion approach, 941nt were deleted, spanning exon 2 (ENSMUSE00000961910, Fig 1C).

All mutant mouse models used were maintained on a C57BL/6NTac background, or on a C57BL/6NTac;Cdh23-repaired background in which the hypomorphic *Cdh23^ahl^* allele present in the C57BL/6 background is corrected to wildtype (*Cdh23^753A>G^*; Mianne, Chessum (27)). No sex-based differences in phenotype were observed during the IMPC adult phenotyping pipeline; as such, both male and female mice were used for all experiments.

### Auditory Brainstem Response (ABR)

Analysis of hearing function by ABR was undertaken as previously described [41]. Briefly, mice were anesthetised with an I.P. injection of ketamine (100 mg ml^-1^ at 10% v/v) and xylazine (20 mg ml^-1^ at 5% v/v) administered at a rate of 0.1 ml/10 g body mass. Animals were placed on a heated mat inside a sound-attenuated chamber (ETS-Lindgren), and electrodes were inserted sub-dermally; below the right pinnae, into the muscle mass below the left ear, and at the cranial vertex. ABR responses were collected, amplified, and averaged using the TDT RZ6 System 3 hardware in conjunction with BioSigRZ (version 5.7.1) software (Tucker Davies Technology). Stimuli were delivered in the form of a 0.1 ms broadband click, or as a single frequency tone at 6, 12, 18, 24, and 30 kHz (5 ms duration, 1 ms rise and fall). All stimuli were presented in 5 dB falling steps from 90 dB SPL, and responses were averaged over 512 (tone) or 300 (click) repeats. Following ABR recordings, mice were either culled by cervical dislocation or recovered with a S.C. injection of atipamezole administered 1 mg kg^-1^ body mass. For the analysis of wave I, supra-threshold traces were further filtered between 400 - 2500Hz to remove additional background noise. Amplitudes were calculated as the difference between the peak and valley. Latencies were calculated as the time from onset to the wave peak.

### Distortion Product Otoacoustic Emissions (DPOAEs)

To assess outer hair cell function in vivo, surgical anaesthesia was induced by intraperitoneal injection of ketamine (100 mg ml^-1^ at 10% v/v), xylazine (20 mg ml^-1^ at 5% v/v) and acepromazine (2 mg ml^-1^ at 8% v/v), administered at a rate of 0.1 ml/10 g body mass. Once the required depth of anaesthesia was confirmed by the lack of the pedal reflex, a section of pinna was removed to allow unobstructed access to the external auditory meatus. Mice were then placed on a heated mat inside a sound-attenuated chamber (ETS-Lindgren) and the DPOAE probe assembly was inserted into the ear canal using a pipette tip to aid correct placement. DPOAE tests were performed using frequency-specific tone-burst stimuli at 6, 12, 18, 24, and 30 kHz with the TDT RZ6 System 3 hardware and BioSigRZ (version 5.7.1) software (Tucker Davis Technology). An ER10B+ low noise probe microphone (Etymotic Research) was used to measure the DPOAE near the tympanic membrane. Tone stimuli were presented via separate MF1 (Tucker Davis Technology) speakers, with f_1_ and f_2_ at a ratio of f_2_/f_1_ = 1.2 (L1 = 65 dB SPL, L2 = 55 dB SPL), centred around the frequencies of 6, 12, 18, 24, and 30 kHz. In-ear calibration was performed before each test. The f1 and f2 tones were presented continuously and a fast-Fourier transform was performed on the averaged response of 356 epochs (each approximately 21 ms). The level of the 2f1 – f2 DPOAE response was recorded and the noise floor was calculated by averaging the four frequency bins on either side of the 2f_1_ – f_2_ frequency. DPOAEs presented as the response above the averaged noise floor. Following DPOAE recordings, mice were culled by cervical dislocation.

### Immunohistochemistry

To assess protein localisation in the cochlea, animals were culled by cervical dislocation and inner ears were removed and fixed by perfusion at the round and oval window with 4% PFA, followed by 1 h submersion fixation on ice. For whole-mounts, ears were finely dissected to expose the sensory epithelium. Sections were cut from whole tissues decalcified for 72 hr at 4°C in 3.5% EDTA, cryoprotected with 30% sucrose, then embedded in OCT before cutting into 12 μm slices on a cryostat. For synaptic labelling, cochleae were permeabilised using PBS + 0.3% Triton X-100, blocked in 5% donkey serum, and incubated at 37°C overnight (16-18 hr) with rabbit anti-Ribeye A domain (1:200, Synaptic Systems Cat# 192 103, RRID:AB_2086775) and mouse anti-GluR2 (1:200, Millipore Cat# MAB397, RRID:AB_2113875). For all other antibodies, cochleae were permeabilised with PBS + 0.1% Triton X-100, blocked in 5% donkey serum + 1% BSA, and incubated with primary antibodies overnight (16-18 hr) at 4°C. Primary antibodies: sheep anti-Np (1:200, R and D Systems Cat# AF7818, RRID:AB_2715517), goat anti-Np65 (1:100, R and D Systems Cat# AF5360, RRID:AB_2155920), mouse anti-PMCA 5F10 (1:200, Thermo Fisher Scientific Cat# MA3-914, RRID:AB_2061566), rabbit anti-PMCA1 (1:200, Alomone Labs Cat# ACP-005, RRID:AB_2756567), rabbit anti-PMCA2 (1:200, Abcam Cat# ab3529, RRID:AB_303878), rabbit anti-Neurofilament 200 (1:200, Sigma-Aldrich Cat# N4142, RRID:AB_477272). To allow the detection of primary antibodies, cochleae were incubated with a relevant fluorophore-conjugated secondary antibody for either 1 hr at room temperature or 2 hr at 37°C (synaptic labelling). Secondary antibodies: AlexaFluor-568 donkey anti-rabbit (1:500, Thermo Fisher Scientific Cat# A10042, RRID:AB_2534017), Alexa Fluor-568 donkey anti-mouse (1:500, Thermo Fisher Scientific Cat# A10037, RRID:AB_2534013), Alexa Fluor-488 donkey anti-rabbit (1:500, Thermo Fisher Scientific Cat# A-21206, RRID:AB_2535792), Alexa Fluor-488 donkey anti-mouse (1:500, Thermo Fisher Scientific Cat# A-21202, RRID:AB_141607), Alexa Fluor-488 donkey anti-sheep (1:500, Thermo Fisher Scientific Cat# A-11015, RRID:AB_2534082), Alexa Fluor-568 donkey anti-goat (1:500, Thermo Fisher Scientific Cat# A-11057, RRID:AB_2534104). Conjugated Phalloidin stains were also used to visualise stereocilia bundles where required: Alexa Fluor-647 Phalloidin (1:200, Thermo Fisher Scientific Cat# A22287), Alexa Fluor-488 Phalloidin (1:200, Thermo Fisher Scientific Cat# A12379). Samples were visualised using a Zeiss LSM 710 with Airyscan detector under either 20x magnification or 63x oil magnification (synapses). Images were processed using the Zeiss Zen microscopy software and Fiji.

### Western Blot

Membrane enriched lysates were extracted from whole cochlea using a Mem-PER plus Membrane Protein Extraction kit (Thermo Scientific^TM^; Cat# 89842). For each genotype, six pooled cochleae (three mice) and a half brain were extracted into separate membrane and cytosolic enriched lysates as per kit instructions. Glycosylation was removed from denatured extracted lysates using PNGase F (NEB, Cat# P0704S) as per the manufacturer’s protocol. Protein (10 mg) was separated with SDS/PAGE and transferred onto 0.45 μm Nitrocellulose membranes (Invitrogen™, Cat# LC2001). Membranes were blocked with 5% BSA for 1 hour at room temperature, then probed with sheep anti-Np (1:1000, R and D Systems Cat# AF7818, RRID:AB_2715517), goat anti-Np65 (1:200, R and D Systems Cat# AF5360, RRID:AB_2155920) overnight (16-18 hr) at 4°C. Equal loading was confirmed with antibodies against β-actin (1:5000, Proteintech Cat# 60008-1-Ig, RRID:AB_2289225), α-tubulin (1:1000, Thermo Fisher Scientific Cat# A11126, RRID:AB_2534135), or Na^2+^/K^+^ ATPase (1:5000, Abcam Cat# ab76020, RRID:AB_1310695). For detection, membranes were incubated with either a HRP-conjugated donkey anti-sheep antibody (1:5000, R and D Systems Cat# HAF016, RRID:AB_562591), and imaged with a UVP ChemiDoc-It imaging system with a BioChemi HR camera; or with a donkey anti-goat IRDye 800CW (1:10000, LI-COR Biosciences Cat# 925-32214, RRID:AB_2687553), a donkey anti-mouse IRDye 800CW (1:10000, LI-COR Biosciences Cat# 926-32212, RRID:AB_621847), or a donkey anti-rabbit IRDye 680RD (1:10000, LI-COR Biosciences Cat# 926-68073, RRID:AB_10954442) and imaged using the Odyssey CLx Infrared Imaging System (LI-COR).

### Single-hair cell electrophysiology

Inner and outer hair cells from *Nptn^tm1b^* mice were studied in acutely dissected organs of Corti from postnatal day 7 (P7) to P25, where the day of birth is P0. Animals were killed by cervical dislocation. Cochleae were dissected in normal extracellular solution (in mM): 135 NaCl, 5.8 KCl, 1.3 CaCl_2_, 0.9 MgCl_2_, 0.7 NaH_2_PO_4_, 5.6 D-glucose, 10 Hepes-NaOH. Sodium pyruvate (2 mM), MEM amino acids solution (50X, without L-Glutamine) and MEM vitamins solution (100X) were added from concentrates (Fisher Scientific, UK). The pH was adjusted to 7.5 (osmolality ~308 mmol kg-1).

Voltage-clamp recordings were performed at room temperature (22-24°C) using an Optopatch amplifier (Cairn Research Ltd, UK). For basolateral membrane current recordings, the patch pipette contained the following intracellular solution (in mM): 131 KCl, 3 MgCl2, 1 EGTA-KOH, 5 Na_2_ATP, 5 Hepes-KOH, 10 Na_2_-phosphocreatine (pH 7.3; osmolality ~296 mmol kg-1). Mechanoelectrical transduction recordings were performed using an intracellular solution containing (in mM): 131 CsCl, 3 MgCl_2_, 1 EGTA-CsOH, 5 Na_2_ATP, 0.3 Na_2_GTP, 5 Hepes-CsOH, 10 Na_2_-phosphocreatine (pH 7.3). Patch pipettes were coated with surf wax. Data acquisition was controlled by pClamp software using a Digidata 1440A board (Molecular Devices, USA). Recordings were low-pass filtered at 2.5 kHz (8-pole Bessel) and sampled at 5 kHz. Data analysis was performed using Origin software (OriginLab, USA). Membrane potentials were corrected for a liquid junction potential measured between electrode and bath solutions, which was −4 mV.

Mechanoelectrical transducer (MET) currents were elicited by stimulating the hair bundles of OHCs using a fluid-jet from a pipette (tip diameter 8-10 μm) driven by a piezoelectric disc as previously described [2, 16, 51]. The pipette tip of the fluid-jet was positioned near to the bundles to elicit a maximal MET current. Mechanical stimuli were applied as force-steps or saturating 50 Hz sinusoids (filtered at 0.25 kHz, 8-pole Bessel) with driving voltages of ± 40 V.

### Experimental Design and Statistical Analyses

Mean values are quoted in text and figures as mean ± standard deviation (SD), except for ABR data which is displayed as median ± interquartile range (IQR). Statistical comparisons of means were made by analysis of variance (one-way or two-way ANOVA) followed by a suitable posttest, or by Student’s two-tailed t-test with or without correction for multiple comparisons. ABR thresholds were second scored by an independent researcher blinded to genotype and overall trends and levels of significance were confirmed to match. For the analysis of ABR thresholds where true values were out of the range of our equipment, thresholds were given a value of 95 dB SPL and marked as “no response” (NR).

## Acknowledgements

The authors wish to thank: J Harrison, and MLC Ward staff for their assistance with mouse husbandry and tamoxifen dosing; and, the Molecular and Cellular Biology group at the Harwell Institute for generating the *Nptn^Δexon2^* mice. This work was supported by: the BBSRC (BB/S006257/1 to W.M. and M.R.B.); the Medical Research Council (MC_UP_1503/2 to M.R.B); and, the Royal National Institute for Deaf People (G81 to S.D.M.B., W.M. and M.R.B.).

